# Multiple Sclerosis Stages and their Differentially Expressed Genes: A Bioinformatics Analysis

**DOI:** 10.1101/2024.01.20.576448

**Authors:** Faten Alaya, Ghada Baraket, Daniel A. Adediran, Katelyn Cuttler, Itunu Ajiboye, Mark T. Kivumbi, Nikita Sitharam, Olaitan I. Awe

## Abstract

Multiple Sclerosis (MS) is an inflammatory, chronic, autoimmune, and demyelinating disease of the central nervous system. MS is a heterogeneous disease with three main clinical forms, affecting the progression and therefore the treatment of the disease. Thus, finding key genes and microRNAs (miRNA) associated with MS stages and analyzing their interactions is important to better understand the molecular mechanism underlying the occurrence and the evolution of MS. Based on publicly available datasets of mRNA and miRNA expression profiles, differentially expressed genes (DEGs) and differentially expressed miRNAs (DEMs) between patients with different stages of MS and healthy controls and between relapsing and remitting phases of RRMS were determined using Deseq2 and GEO2R tools. We then analyzed miRNA-mRNA regulatory interactions and gene ontology for the DEGs.

Based on miRNA-mRNA regulatory interactions, we identified potential biomarkers of RRMS, 13 upregulated miRNA regulators of 30 downregulated genes and 17 downregulated miRNA regulators of 32 upregulated genes. We also identified 9 downregulated miRNA regulators of 12 upregulated genes as potential biomarkers of SPMS.

Our study findings highlight some key protein-coding genes and miRNAs that are involved in the occurrence and evolution of MS.

## Introduction

Multiple sclerosis (MS) is an inflammatory, autoimmune, demyelinating disease of the central nervous system (CNS) (Sospedra and Martin, 2016). It is the most common and leading cause of severe disability of non-traumatic origin in people in their thirties. It affects nearly 2.8 million people worldwide with an average sex ratio of 3 to 1 (women: men) and an average age of onset of 32 years (mapping multiple sclerosis around the world). The exact cause of MS is not yet known. Rather, it has a multifactorial origin, consisting of environmental and genetic factors (Jackle *et al*., 2020). In addition to this, there are viral infections during childhood which can play an important role in the development of the disease and also in triggering the autoimmune process (Dyment *et al*., 2004).

The main target of MS is the myelin sheath (Safari-Alighiarloo *et al*., 2020). It is characterized by the appearance of demyelinating lesions that are inflammatory in nature (Karussis, 2014). These are generally in the white matter and distributed throughout the myelinated areas of the CNS, which prevents the conduction of nerve impulses and explains the appearance of various clinical signs (Baecher-Allan *et al*., 2018). In the later stages of multiple sclerosis, axonal damage also occurs (Trapp *et al*., 1998). In a genetically susceptible subject, a multi-step immuno-pathological process involving an autoreactive lymphocyte clone recognizing myelin as an autoantigen is responsible for triggering autoimmune attacks and the development of MS (Freiesleben *et al*., 2016). Since MS can affect several areas of the CNS, multiple different symptoms can be observed. However, common symptoms can include fatigue, weakness, sensory loss and ataxia (Ben-Zacharia, 2011). MS is an extremely heterogeneous disease which manifests differently, both in the expression of symptoms and in the way it evolves, depending on the stage and the clinical form of the disease (Tullman, 2013).

There are three clinical forms of MS: Relapsing-Remitting MS (RRMS), Secondary Progressive MS (SPMS) and Primary Progressive MS (PPMS) (Klineova and Lublin, 2018) (Fig. 1). RRMS is the most common form of MS (85% of cases) and is characterized by relapses which reach a plateau over several weeks and then gradual recovery (Fig. 1A). In early MS, complete recovery can be seen. However, as the disease develops, most of the time there is incomplete recovery and remaining damage. In addition, for every clinical recovery, there are so-called ‘asymptomatic’ lesions which can lead to future relapses. SPMS generally develops 10-15 years after RRMS onset, and it gradually progresses from faster distinct relapses to slowly progressive disease (Fig. 1B). However, in 10-15% of cases, an individual presents with PPMS. Individuals in this stage of the disease manifest a permanent progression from the onset of MS, typically involving one dominant neuronal system (Fig. 1C) (Dobson and Giovannoni, 2019).

**Figure 1.**
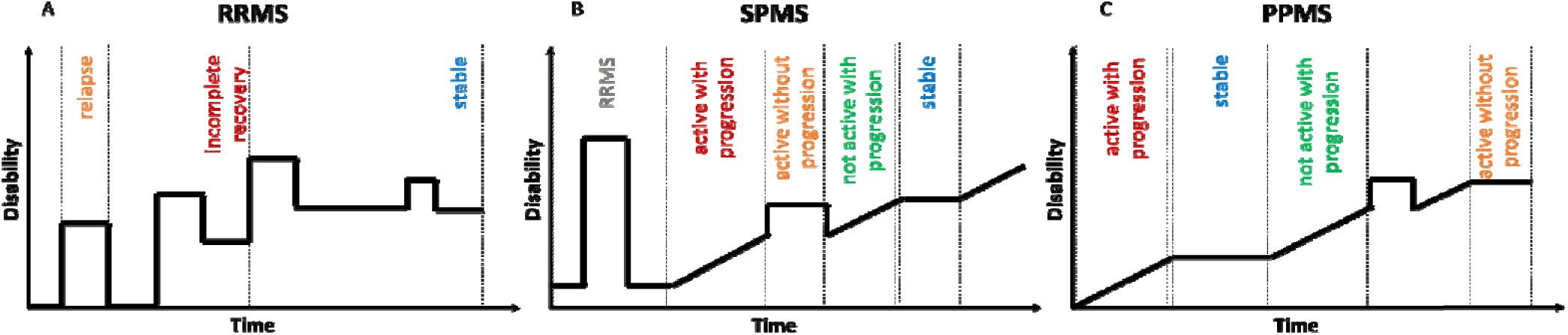
Different stages of multiple sclerosis <u>(Lublin</u> *et al*., <u>2014)</u>.

**Figure 2.**
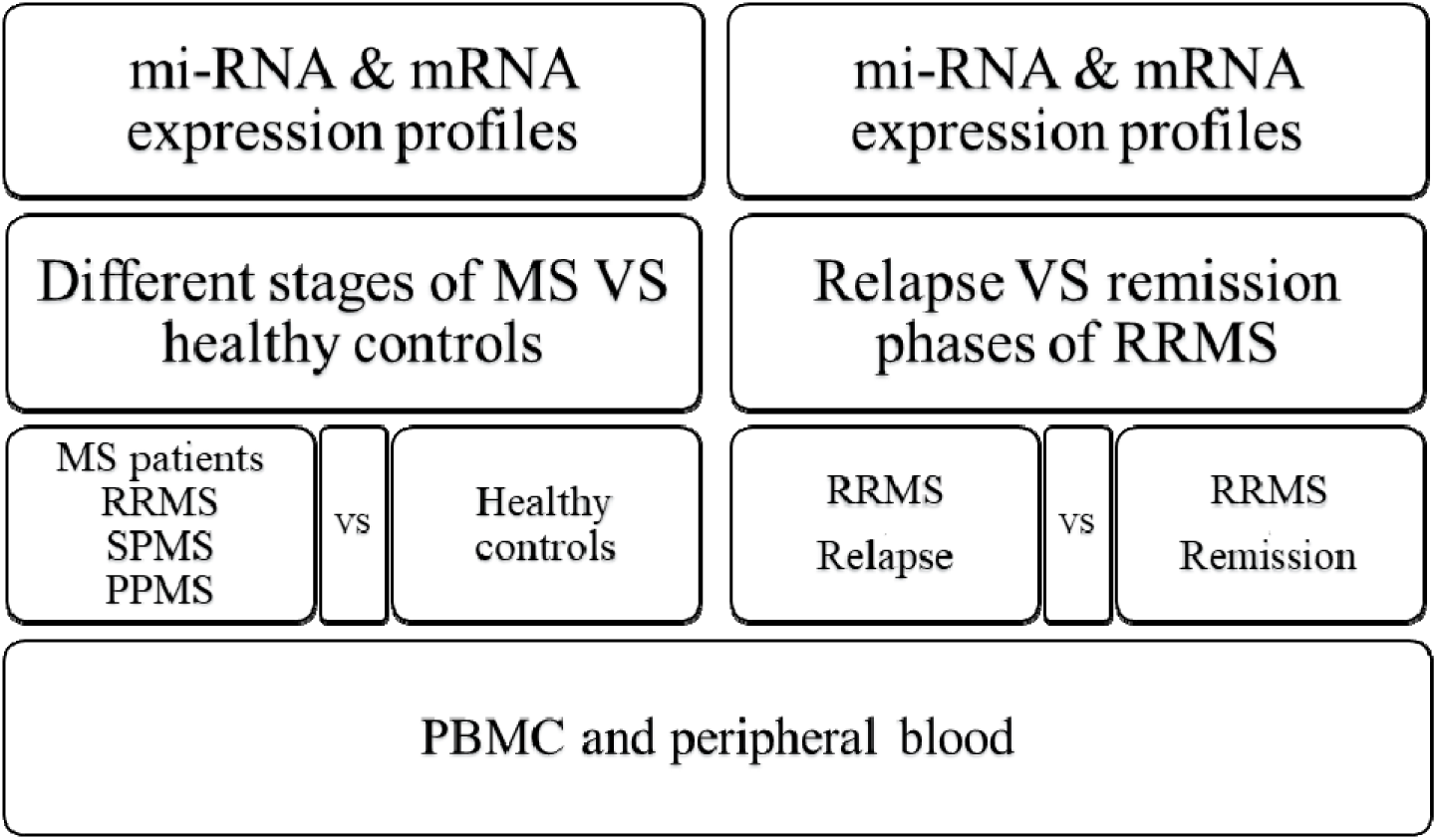
Data extracted from the GEO database.

**Figure 3.**
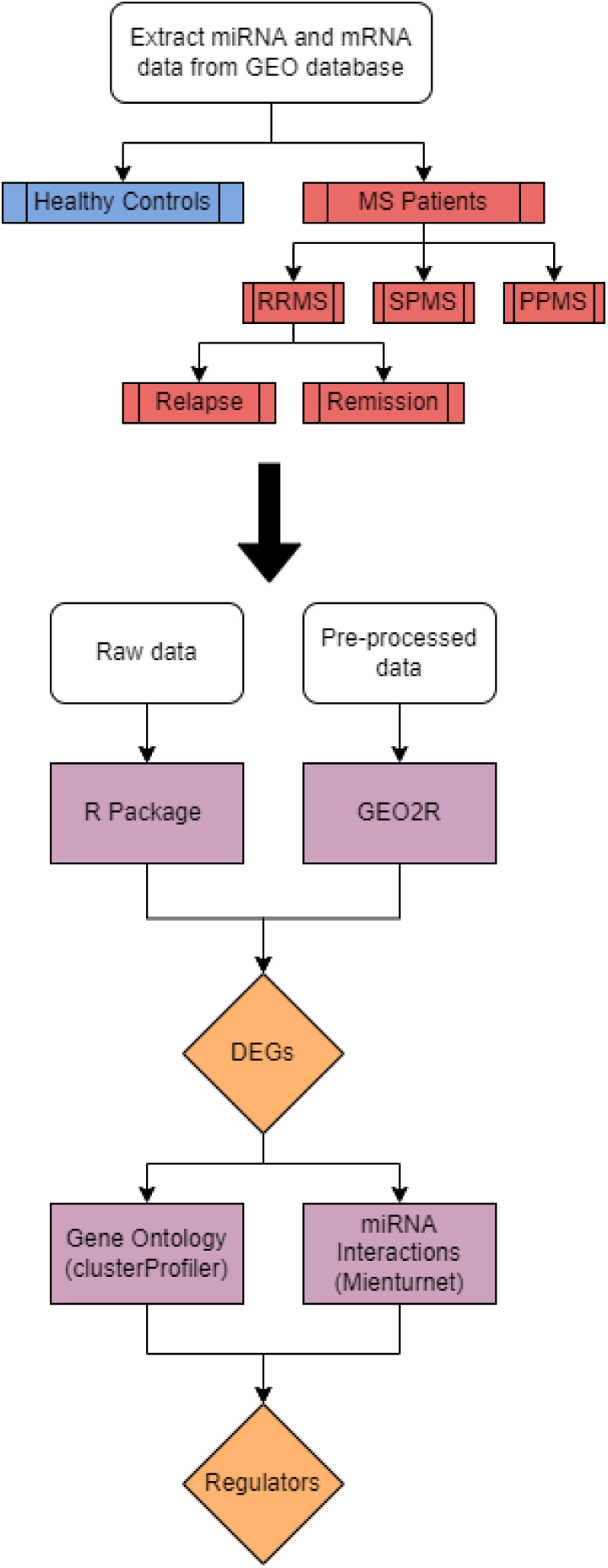
Workflow followed in the study.

Determining the clinical form of MS in a patient is usually done after the diagnosis following several examinations performed by physicians. Therefore, it is very time-consuming and expensive. However, the early determination of the clinical form of the disease and the possible prediction of its evolution can help to improve its management. Therefore, there is a vital need to find biological markers (or biomarkers) that can distinguish the stages of the disease (Klineova and Lublin, 2018; Mayeux, 2004). In addition, recent studies have shown that there are indeed genetic differences and specific differential gene expression pattern differences between RRMS and PPMS (Jia *et al*., 2018; Koch *et al*., 2018). In other clinical applications, mutations in specific genes can also be used to screen for disorders in newborns (Wesonga and Awe, 2022) and as sequencing technology develops, genomics will rely heavily on our ability to use sequencing data for applications like biomarker discovery, viral pathogen evolution (Awe *et al*., 2023; Oluwagbemi *et al*., 2018), bioinformatics workflow development (Ather *et al*., 2018) or gene expression in plants (Die *et al*., 2019). Recently, it has been demonstrated that the aberrant expression of microRNAs (miRNAs) is involved in the pathogenesis of many diseases and that they could be potential biomarkers of many disorders including MS (O’Brien *et al*., 2018; Waschbisch *et al*., 2011). miRNAs are a class of small, single-stranded, non-coding RNAs that act as post-transcriptional regulators of gene expression (Housley *et al*., 2015). When paired with a target mRNA, miRNAs can lead to the inhibition of its translation or its degradation (Ridolfi *et al*., 2013).

Moreover, advancements in genomics and molecular biology have opened avenues for exploring the intricate genetic landscape of diseases. Nyamari *et al*. (2023) delved into the expression level analysis of the ACE2 receptor gene in African-American and non-African-American COVID-19 patients, shedding light on genetic factors influencing susceptibility to the virus. Nzungize *et al*. (2022) conducted transcriptional profile analysis of COVID-19 and malaria patients, while El Abed *et al*., (2023) conducted transcriptional profile analysis revealing potential biomarkers in epilepsy patients, and emphasizing the broader applications of gene expression studies in understanding various diseases. Additionally, Obura *et al*. (2022) explored the molecular phylogenetics of HIV-1 subtypes in African populations, providing insights into the genetic diversity of the virus in Sub-Saharan African countries. Furthermore, Ogbodo *et al*. (2023) applied computational methods for identifying potential inhibitors targeting cdk1 in colorectal cancer, showcasing the integration of genomics and computational approaches in cancer research. In the realm of infectious diseases, Mwanga *et al*. (2023) developed an enhanced deep convolutional neural network for SARS-CoV-2 variants classification, highlighting the pivotal role of artificial intelligence in understanding and combating emerging viral strains.

Usually, the comparison of miRNA expression is made between MS patients and healthy controls, without separating the stages of the disease (Sohan Forooshan Moghadam *et al*., 2020). A few studies have studied the relapsing-remitting phases of RRMS (Irizar *et al*., 2014; Keller *et al*., 2009). However, very few studies have compared patients belonging to different stages of the disease with healthy controls (Cox *et al*., 2010).

The objectives of this study are therefore i) to analyze available data from various populations, ii) determine the differential expression of mRNA genes and miRNAs between patients with different stages of MS and healthy controls, iii) determine the differential expression of genes and miRNAs between both relapsing and remitting phases of RRMS, and iv) to identify new biomarker (miRNA) associated with MS stages.

## Methods

### 1. Data collection

The NCBI Gene Expression Omnibus (GEO, http://www.ncbi.nlm.nih.gov/geo) (Barrett *et al*., 2013) was queried in order to extract public data using the keyword “multiple sclerosis”. Then, we applied the following filters to the search results: “Expression profiling by array” and “Homo Sapiens”. miRNA and mRNA expression profiles of patients in different stages of MS, patients with both relapsing and remitting phases of RRMS and healthy controls were used.

### 1.1 Comparison between patients with different stages of MS and healthy controls

The following data was used for comparison between patients with different stages of MS and healthy controls: GSE124900 (Baulina *et al*., 2019a; Baulina *et al*., 2019b), GSE21079 (Mathew B Cox *et al*., 2010a; Mathew B Cox *et al*., 2010b), GSE17048 (Booth and Gandhi, 2010; Gandhi *et al*., 2010; Riveros *et al*., 2010) and GSE41890 (Edgar, Domrachev, and Lash, 2002; H Irizar and Otaegui, 2013). Details of the datasets are provided in Table 1 of the supplementary section.

**Table 1:**
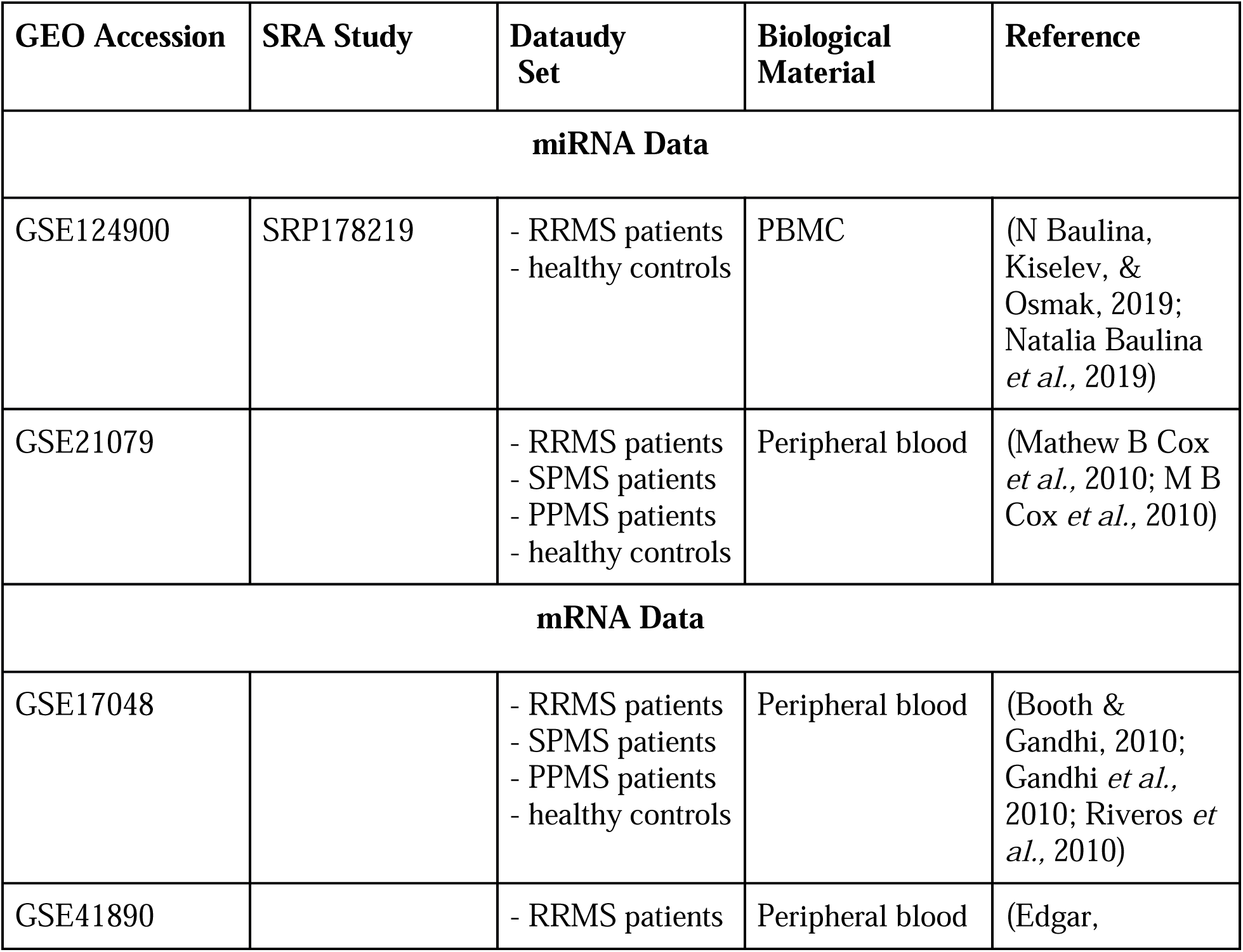
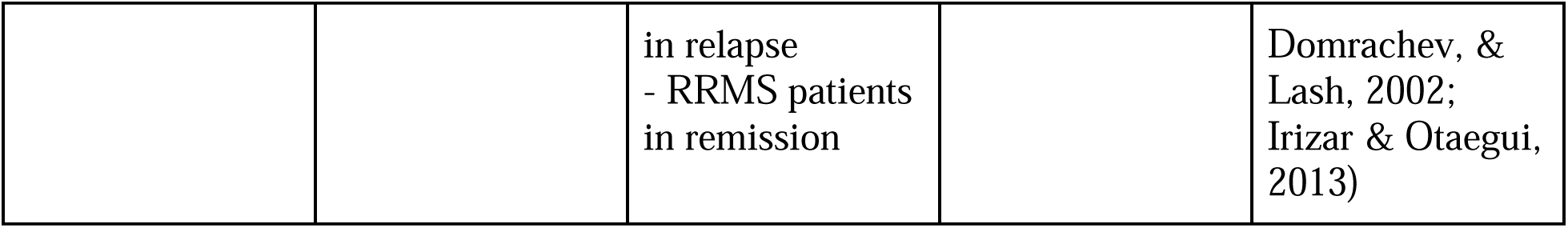
Data sets included in the study.

**Table 2:** miRNA-mRNA regulatory relation from GSE124900 and GSE41890.

| miRNA | miRNA<br>expression | Gene target | Gene<br>expression | Gene<br>description |
| --- | --- | --- | --- | --- |
| **Women: Remission (RRMS-rem) vs Healthy controls (HC)** |  |  |  |  |
| hsa-miR-142-5p | Down-regulated | EGR3 | Up-regulated | early growth<br>response 3 |
|  |  | EGR2 |  | early growth<br>response 2 |
| hsa-miR-153<br>(hsa-miR-153-3p) | Down-regulated | CACNA1C | Up-regulated | calcium voltage-gated channel<br>subunit alpha1 C |
| hsa-miR-92a<br>(hsa-miR-92a-3p) | Up-regulated | SOX4 | Down-regulated | SRY-box 4 |
|  |  | GDF11 |  | growth<br>differentiation<br>factor 11 |
|  |  | TFDP2 |  | transcription<br>factor Dp-2 |
|  |  | EOMES |  | eomesodermin |
|  |  | PLEKHA1 |  | pleckstrin<br>homology<br>domain<br>containing A1 |
|  |  | SLC4A10 |  | solute carrier<br>family 4 member<br>10 |
|  |  | SLC12A2 |  | solute carrier<br>family 12<br>member 2 |
|  |  | DLG5 |  | discs large<br>MAGUK<br>scaffold protein 5 |
|  |  | CRTAM |  | cytotoxic and<br>regulatory T-cell<br>molecule |
|  |  | PHLDA1 |  | pleckstrin<br>homology like<br>domain family A<br>member 1 |
| hsa-miR-141<br>(hsa-miR-141-3p) | Down-regulated | EGR2 | Up-regulated | early growth<br>response 2 |
| hsa-miR-27a<br>(hsa-miR-27a-3p) | Down-regulated | KIAA0319L | Up-regulated | KIAA0319 like |
|  |  | EGR3 |  | EGR3 |
| hsa-miR-374b*<br>(hsa-miR-374b-5p) | Down-regulated | SLC7A14 | Up-regulated | solute carrier<br>family 7 member<br>14 |
|  |  | EGR2 |  | early growth<br>response 2 |
| Men: Relapse (RRMS-rel) vs Healthy controls (HC) |  |  |  |  |
| hsa-miR-376c<br>(hsa-miR-376c-3p) | Up-regulated | IGSF9B | Down-regulated | immunoglobulin<br>superfamily<br>member 9B |
| hsa-miR-532-5p | Down-regulated | GBP1 | Up-regulated | guanylate<br>binding protein 1 |
|  |  | GBP3 |  | guanylate<br>binding protein 3 |
| hsa-miR-409-3p | Up-regulated | BACH2 | Down-regulated | BTB domain and<br>CNC homolog 2 |
|  |  | SCN1A |  | sodium voltage-gated channel<br>alpha subunit 1 |
| hsa-miR-136*<br>(hsa-miR-136-5p) | Up-regulated | LIFR | Down-regulated | LIF receptor alpha |
|  |  | RGS4 |  | regulator of G protein signalling 4 |
| hsa-miR-19a<br>(hsa-miR-19a-3p) | Up-regulated | EBF1 | Down-regulated | early B-cell factor 1 |
|  |  | CSRNP3 |  | cysteine and serine-rich nuclear protein 3 |
|  |  | COL19A1 |  | collagen type XIX alpha 1 chain |
| hsa-miR-758<br>(hsa-miR-758-3p) | Up-regulated | BACH2 | Down-regulated | BTB domain and CNC homolog 2 |
|  |  | OSBPL10 |  | oxysterol binding protein-like 10 |
|  |  | SCN1A |  | sodium voltage-gated channel alpha subunit 1 |
| hsa-miR-377<br>(hsa-miR-377-3p) | Up-regulated | CSRNP3 | Down-regulated | cysteine and serine-rich nuclear protein 3 |
|  |  | CFTR |  | cystic fibrosis transmembrane conductance regulator |
| hsa-miR-485-5p | Up-regulated | CELSR2 | Down-regulated | cadherin EGF LAG seven-pass G-type receptor 2 |
|  |  | CELF3 |  | CUGBP Elav-like family member 3 |
|  |  | OSBPL10 |  | oxysterol binding protein-like 10 |
| hsa-miR-1 (hsa-miR-1-3p) | Down-regulated | TMSB4X | Up-regulated | thymosin beta 4, X-linked |
|  |  | ABCA1 |  | ATP binding cassette subfamily A member 1 |
|  |  | CRISPLD2 |  | cysteine-rich secretory protein LCCL domain containing 2 |
| hsa-miR-493* (hsa-miR-493-5p) | Up-regulated | SYVN1 | Down-regulated | synoviolin 1 |
|  |  | CTNND2 |  | catenin delta 2 |
|  |  | SERTAD2 |  | SERTA domain containing 2 |
|  |  | CSRNP3 |  | cysteine and serine-rich nuclear protein 3 |
|  |  | GPR37L1 |  | G protein-coupled receptor 37 like 1 |
|  |  | ADARB2 |  | adenosine deaminase, RNA-specific B2 (inactive) |
|  |  | SCN1A |  | sodium voltage-gated channel alpha subunit 1 |
|  |  | PDPK1 |  | 3-phosphoinositide-dependent protein kinase 1 |
|  |  | IGSF9B |  | immunoglobulin superfamily member 9B |
| hsa-miR-381<br>(hsa-miR-381-3p) | Up-regulated | SYVN1 | Down-regulated | synoviolin 1 |
|  |  | EBF1 |  | early B-cell factor 1 |
|  |  | CSRNP3 |  | cysteine and serine-rich nuclear protein 3 |
|  |  | OSBPL10 |  | oxysterol binding protein-like 10 |
|  |  | NCAM2 |  | neural cell adhesion molecule 2 |
|  |  | CFTR |  | cystic fibrosis transmembrane conductance regulator |
| hsa-miR-146b-5p | Down-regulated | ACYP2 | Up-regulated | acylphosphatase 2 |
| hsa-miR-485-3p | Up-regulated | BACH2 | Down-regulated | BTB domain and CNC homolog 2 |
|  |  | WIF1 |  | WNT inhibitory factor 1 |
| hsa-miR-30e*<br>(hsa-miR-30e-5p) | Down-regulated | ACVR1 | Up-regulated | activin A receptor type 1 |
|  |  | GBP2 |  | guanylate binding protein 2 |
|  |  | SEC24A |  | SEC24 homolog A, COPII coat complex component |
| hsa-miR-19b<br>(hsa-miR-19b-3p) | Up-regulated | EBF1 | Down-regulated | early B-cell factor 1 |
|  |  | CSRNP3 |  | cysteine and serine-rich nuclear protein 3 |
|  |  | COL19A1 |  | collagen type XIX alpha 1 chain |
| hsa-miR-495<br>(hsa-miR-495-3-p) | Up-regulated | CTNND2 | Down-regulated | catenin delta 2 |
|  |  | ANKRD7 |  | ankyrin repeat domain 7 |
|  |  | GPR37L1 |  | G protein-coupled receptor 37 like 1 |

**Table 3:** miRNA-mRNA regulatory relation from GSE21079 and GSE17048.

| miRNA | miRNA expression | Gene target | Gene expression | Gene description |
| --- | --- | --- | --- | --- |
| SPMS vs HC |  |  |  |  |
| hsa-let-7f, i,d,g | downregulated | ZBTB16 | Upregulated | zinc finger and BTB domain containing 16 |
|  |  | DLST |  | dihydrolipoamide S-succinyltransferase |
|  |  | NAA20 |  | N(alpha)-acetyltransferase 20, NatB catalytic subunit |
| hsa-miR-,15a,15b | downregulated | GIMAP7 | upregulated | GTPase, IMAP family member 7 |
|  |  | ZBTB16 |  | zinc finger and BTB domain containing 16 |
|  |  | DLST |  | dihydrolipoamide S-succinyltransferase |
|  |  | CD69 |  | CD69 molecule |
|  |  | DPH5 |  | diphthamide biosynthesis 5 |
| hsa-miR-20a,20b | downregulated | SNX8 | upregulated | sorting nexin 8 |
|  |  | CD69 |  | CD69 molecule |
| hsa-miR-96 | downregulated | GCNT1 | upregulated | glucosaminyl (N-acetyl) transferase 1, core 2 |
|  |  | MFHAS1 |  | malignant fibrous histiocytoma amplified sequence 1 |
| hsa-miR-148b | downregulated | SNX8 | upregulated | sorting nexin 8 |
|  |  | ZBTB16 |  | zinc finger and BTB domain containing 16 |
| hsa-miR-26b | downregulated | ZBTB16 | upregulated | zinc finger and BTB domain containing 16 |
|  |  | METAP2 |  | methionyl aminopeptidase 2 |
|  |  | HINT1 |  | histidine triad nucleotide-binding protein 1 |
|  |  | NDUFS4 |  | NADH: ubiquinone oxidoreductase subunit S4 |
|  |  | LPAR6 |  | lysophosphatidic acid receptor6 |
| RRMS vs HC |  |  |  |  |
| hsa-miR-20a | downregulated | ZNF652 | upregulated | zinc finger protein 652 |
|  |  | PPP3R1 |  | protein phosphatase 3 regulatory subunit B, alpha |
|  |  | LZIC |  | leucine zipper and CTNNBIP1 domain containing |
|  |  | DUSP8 |  | dual specificity phosphatase 8 |
|  |  | PPP6R2 |  | protein phosphatase 6 regulatory subunit 2 |
|  |  | RPS6KA3 |  | ribosomal protein S6 kinase A3 |
|  |  | RAB12 |  | RAB12, member RAS oncogene family |
|  |  | GBF1 |  | Golgi brefeldin A resistant guanine nucleotide exchange factor 1 |
| hsa-let-7g hsa-miR-98,hsa-let-7f, | downregulated | ZF652 | upregulated | zinc finger protein 652 |
|  |  | DLST |  | dihydrolipoamide S-succinyltransferase |
|  |  | RPS6KA3 |  | ribosomal protein S6 kinase A3 |
|  |  | CELF1 |  | CUGBP, Elav-like family member 1 |
|  |  | L2HGDH |  | L-2- |
|  |  |  |  | hydroxyglutarate dehydrogenase |
|  |  | LZIC |  | leucine zipper and CTNNBIP1 domain containing |
| hsa-miR-93 | downregulated | SUGT1 | upregulated | SGT1 homolog, MIS12 kinetochore complex assembly cochaperone |
|  |  | RAB12 |  | RAB12, member RAS oncogene family |
|  |  | PPP6R2 |  | protein phosphatase 6 regulatory subunit 2 |
| hsa-miR-15b | downregulated | GIMAP7 | upregulated | GTPase, IMAP family member 7 |
|  |  | ZBTB16 |  | zinc finger and BTB domain containing 16 |
|  |  | DLST |  | dihydrolipoamide S-succinyltransferase |
|  |  | CD69 |  | CD69 molecule |
|  |  | DPH5 |  | diphthamide biosynthesis 5 |

**Table 4:**
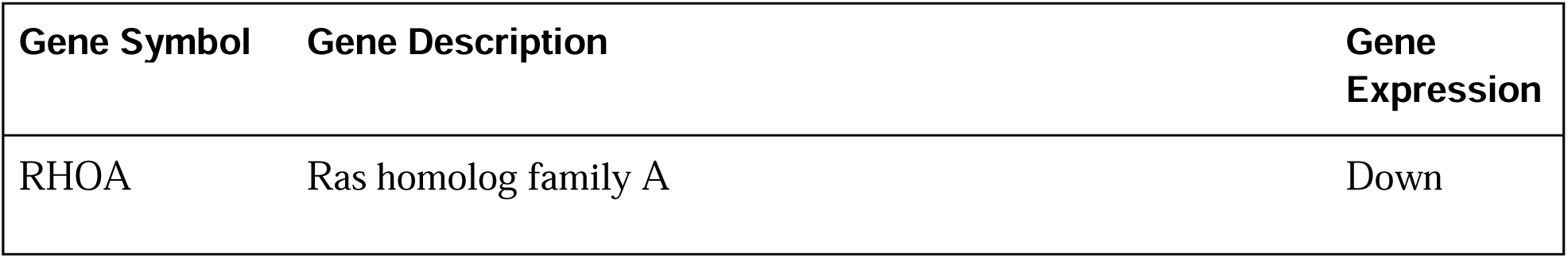
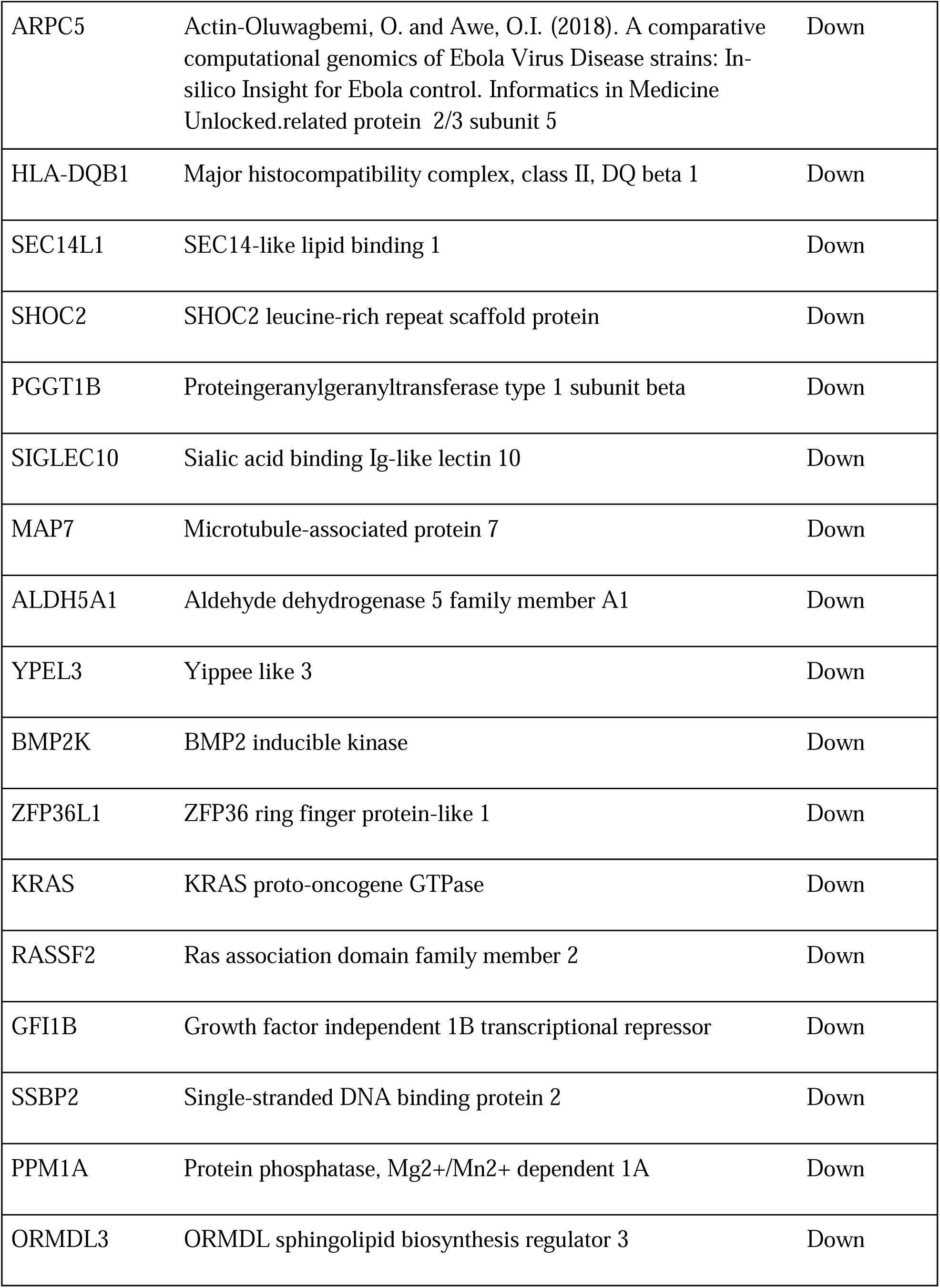
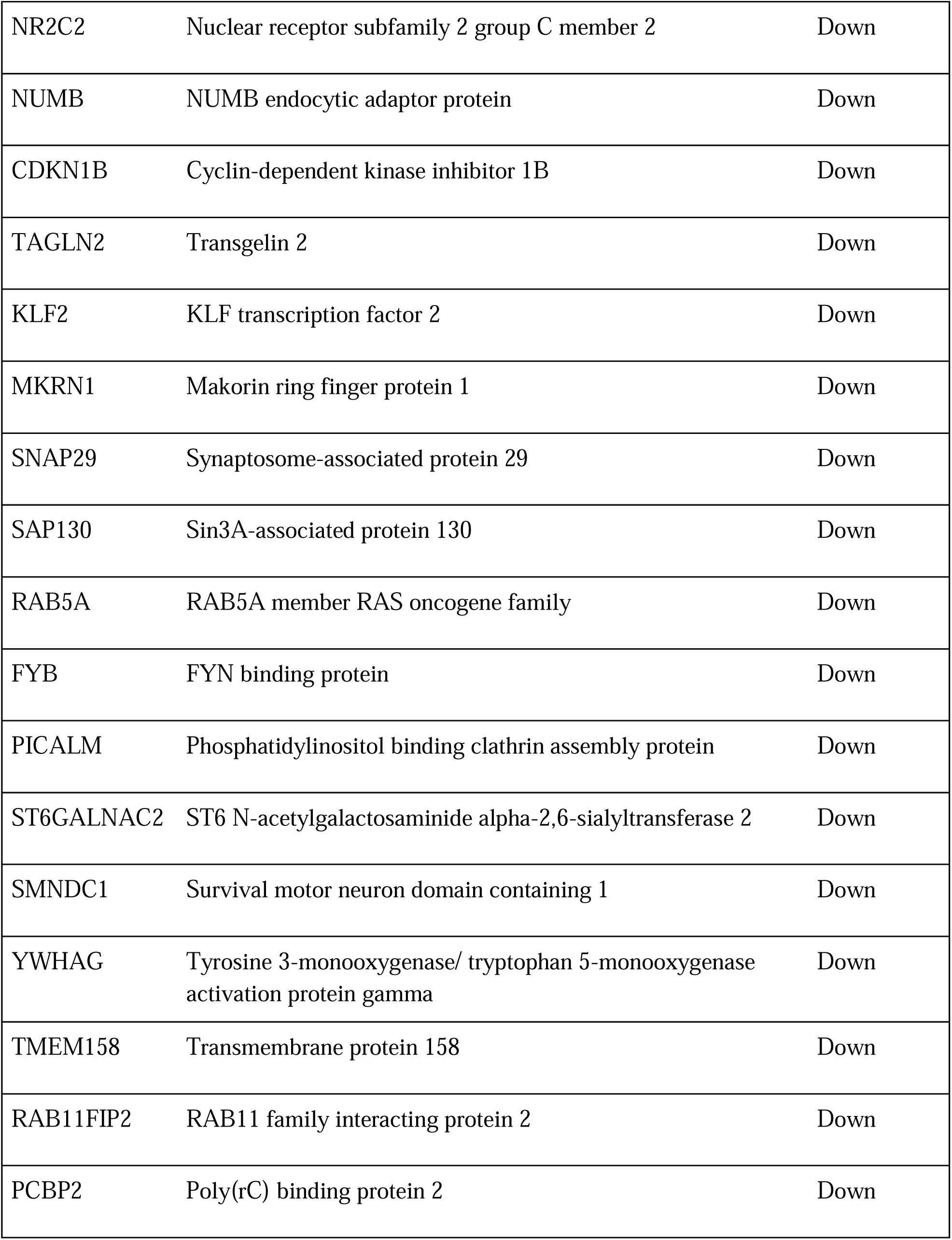
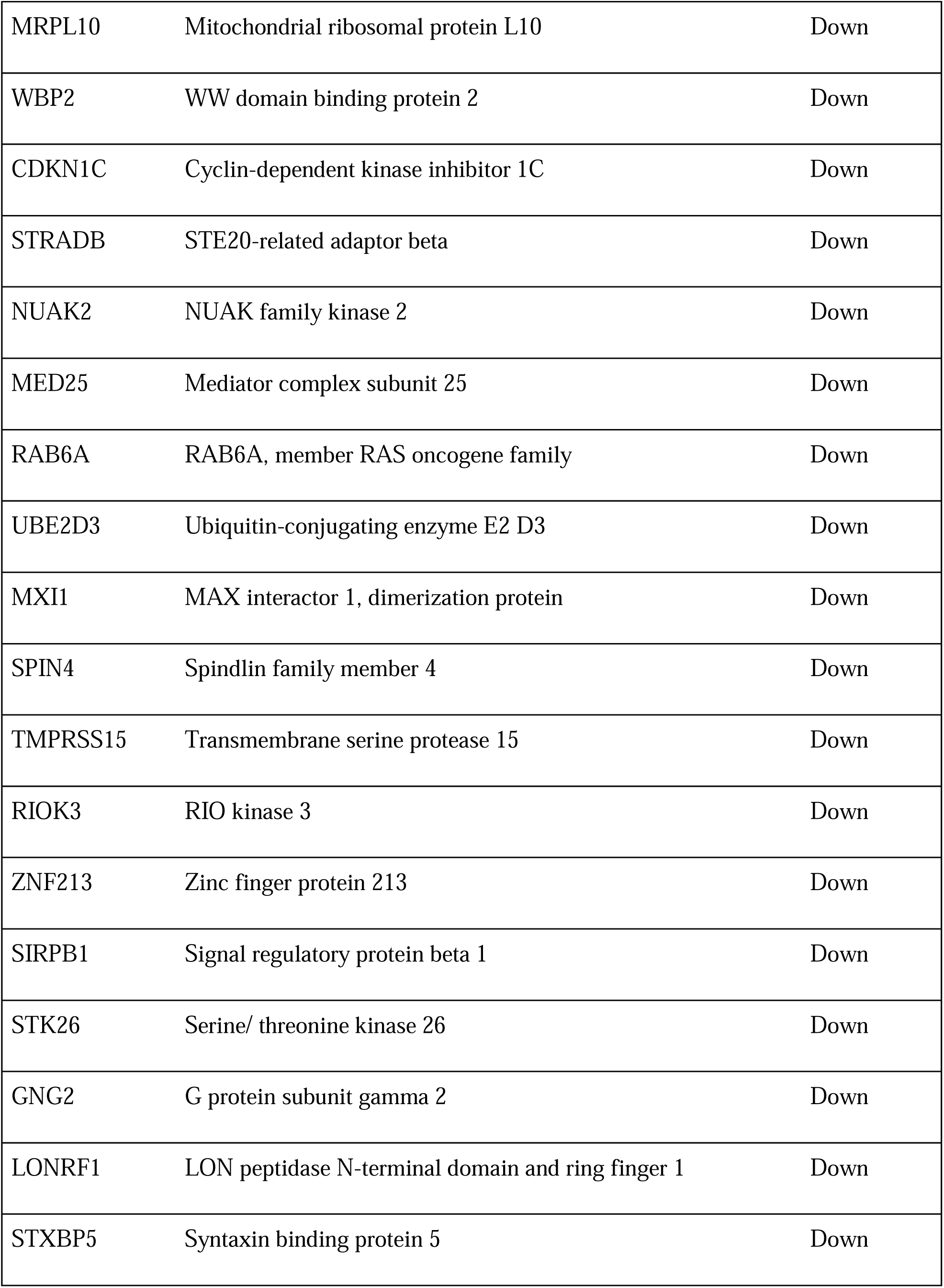
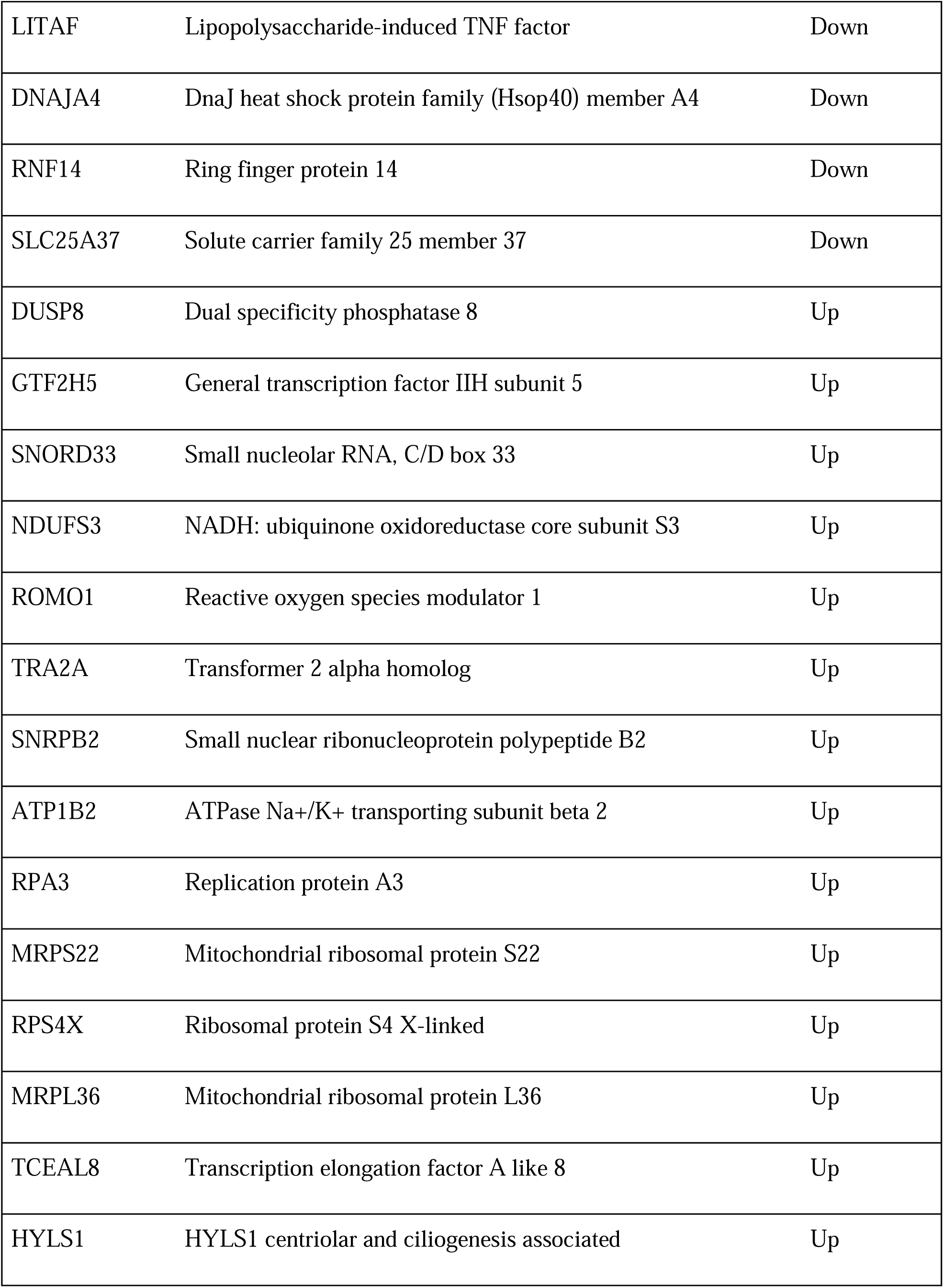
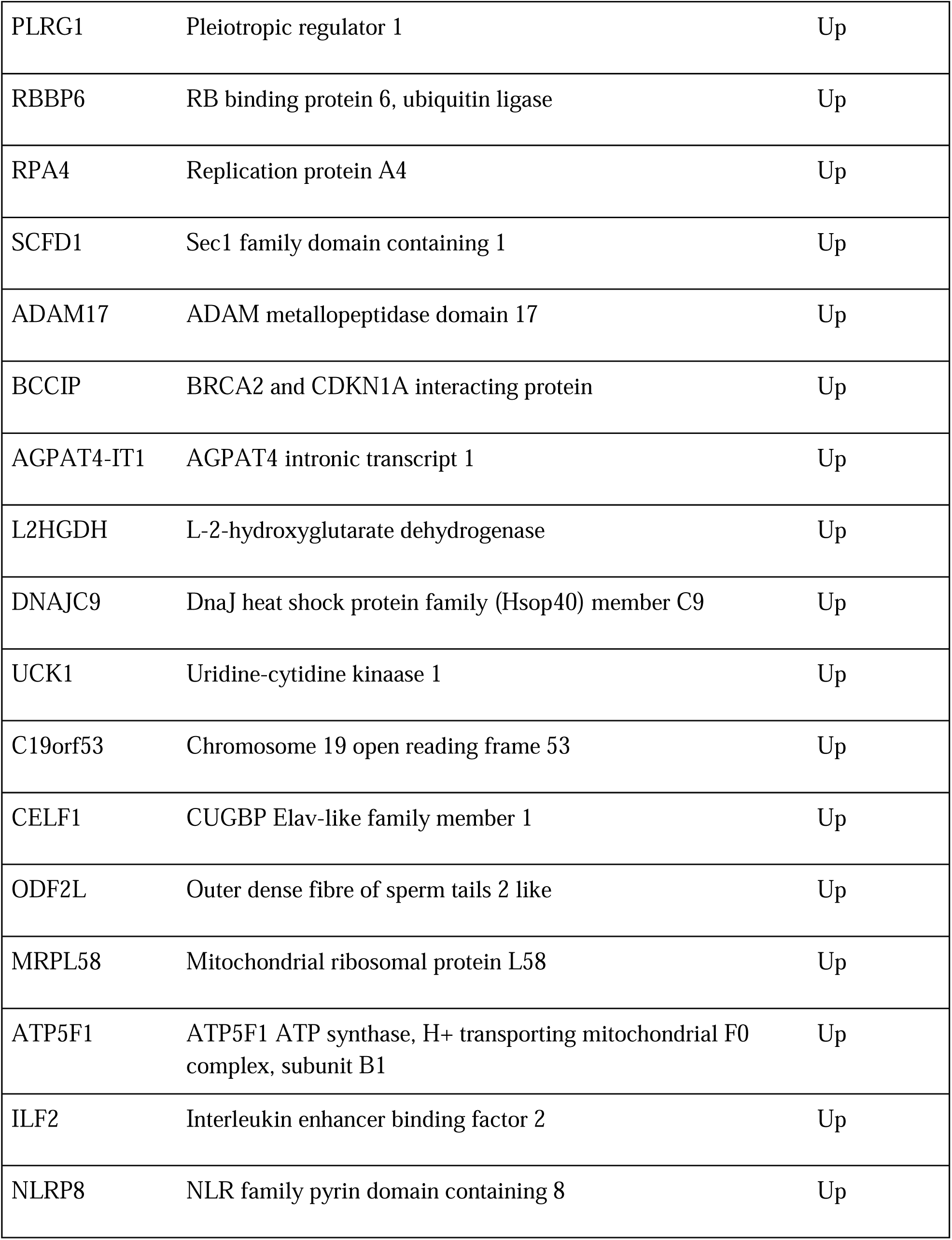
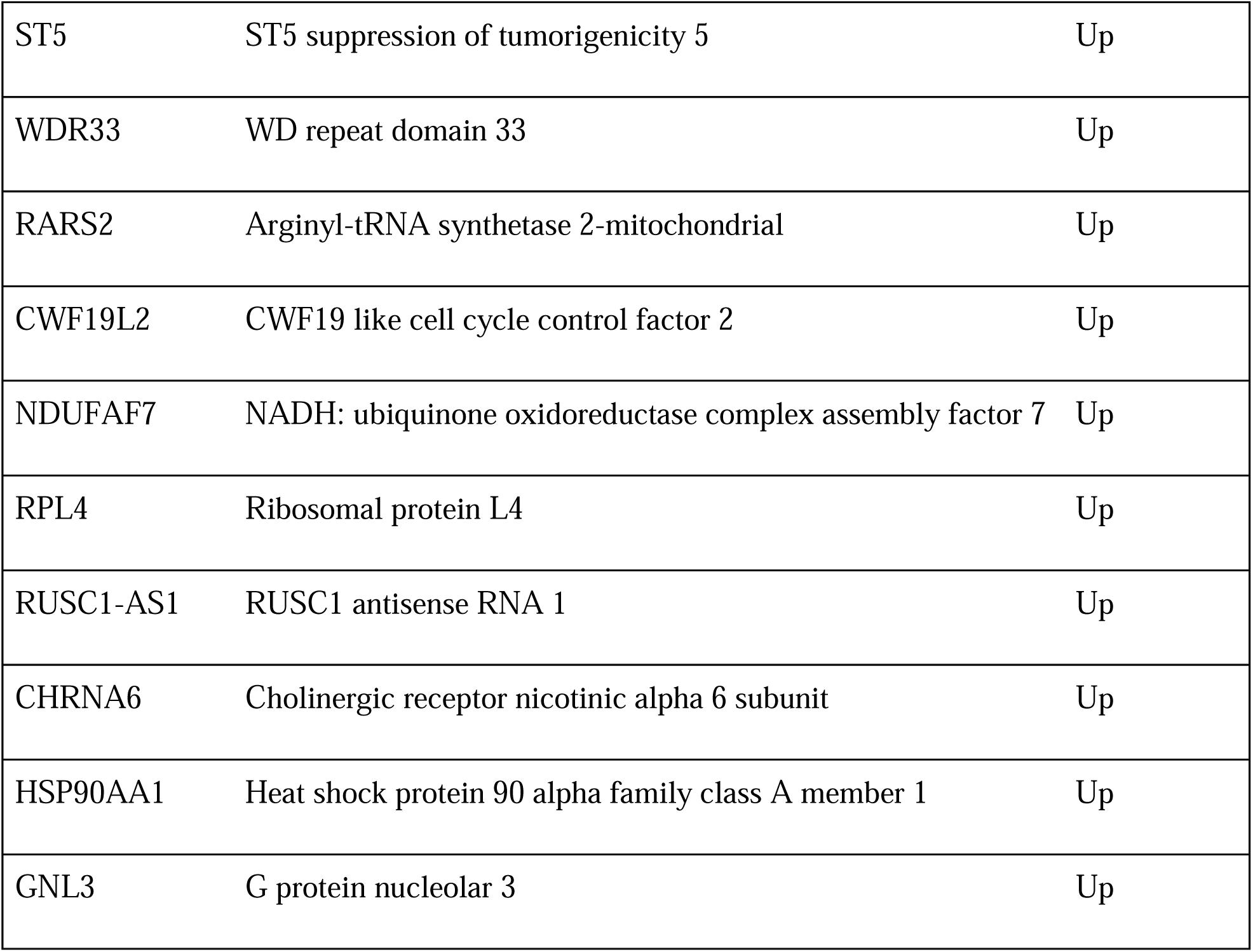
Common mRNAs between GSE17048 and GSE41890.

**Table 5:**
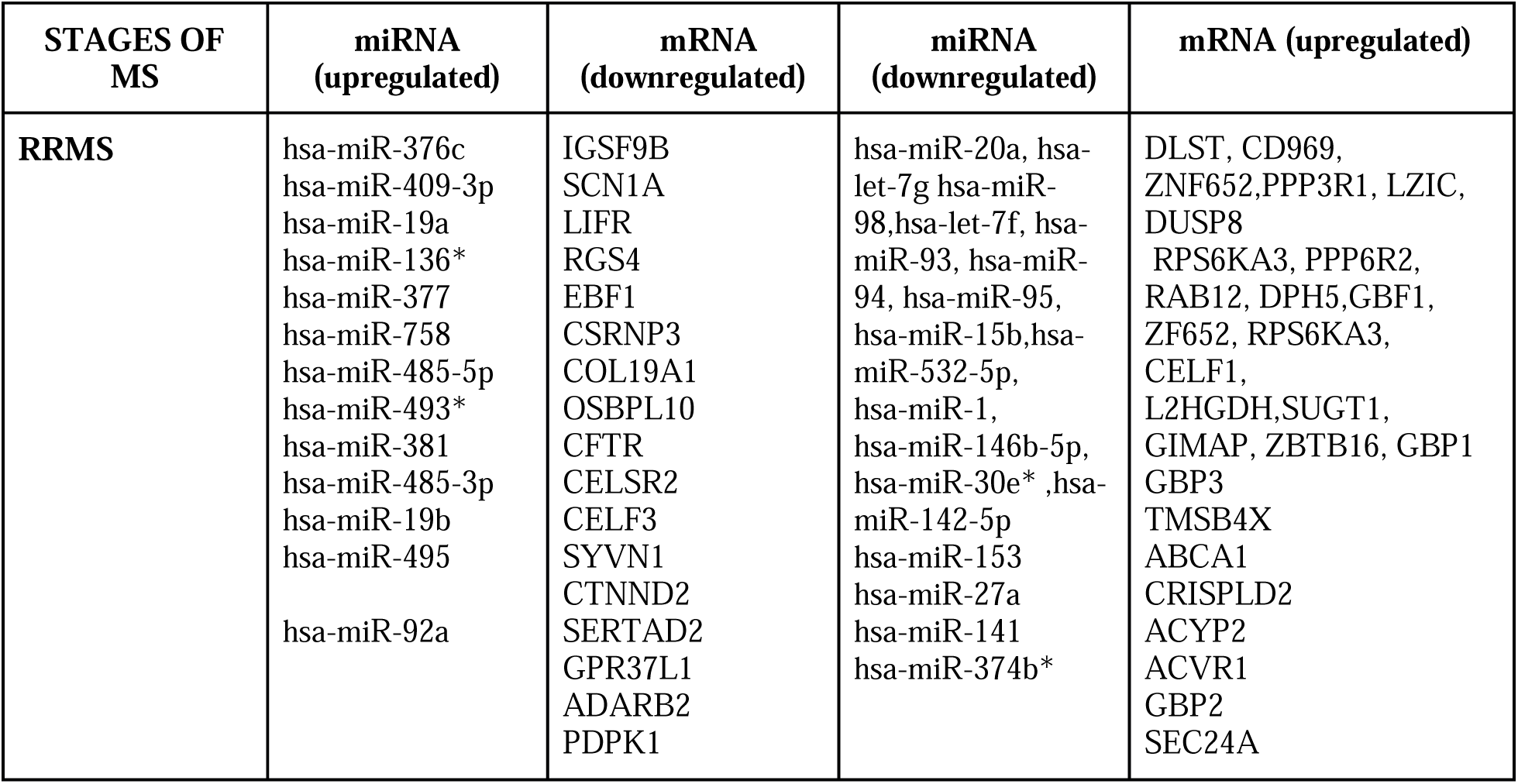
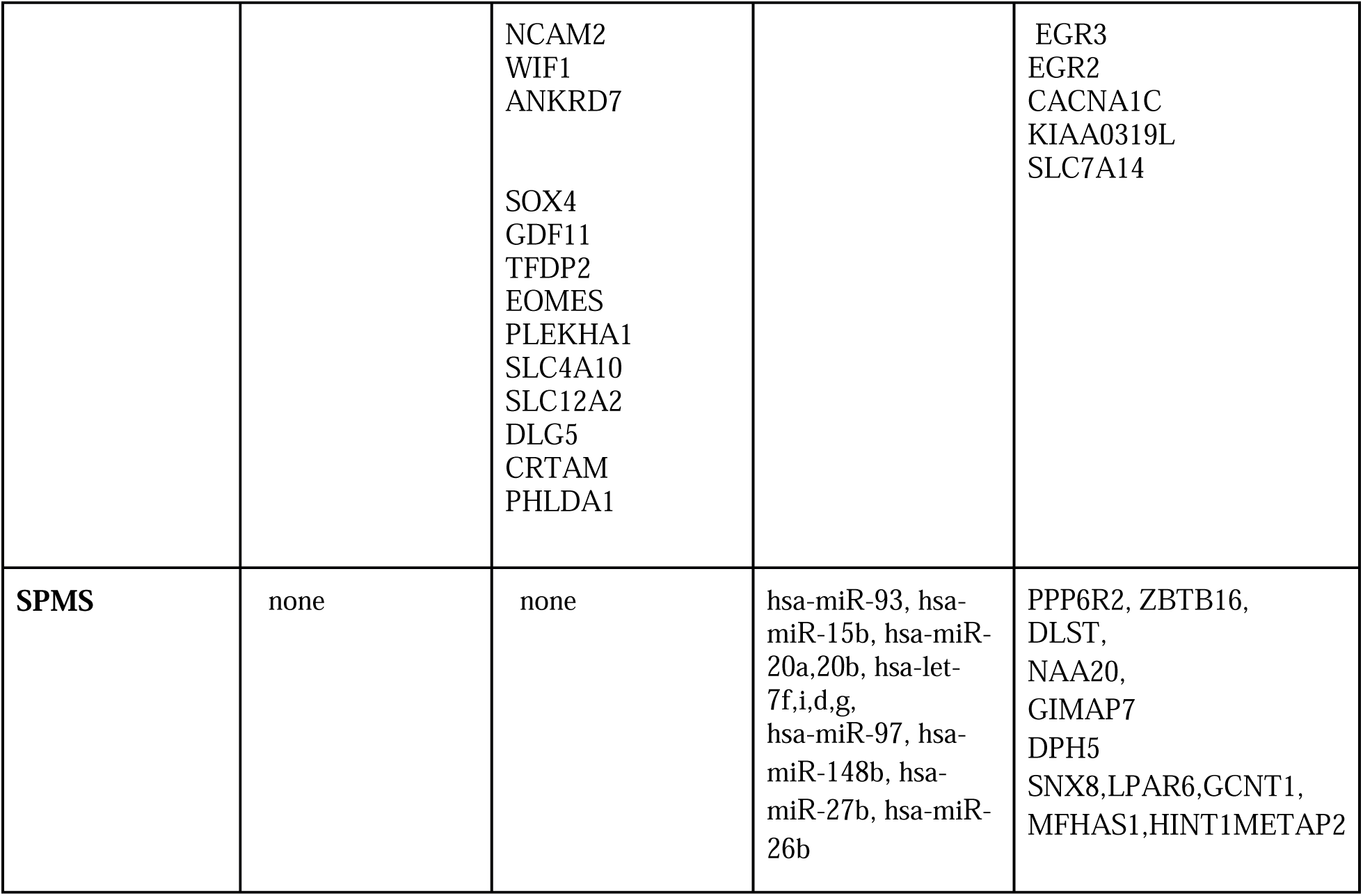
Biomarker discovery.

| STAGES OF MS | miRNA (upregulated) | mRNA (downregulated) | miRNA (downregulated) | mRNA (upregulated) |
| --- | --- | --- | --- | --- |
| **RRMS** | hsa-miR-376c<br>hsa-miR-409-3p<br>hsa-miR-19a<br>hsa-miR-136*<br>hsa-miR-377<br>hsa-miR-758<br>hsa-miR-485-5p<br>hsa-miR-493*<br>hsa-miR-381<br>hsa-miR-485-3p<br>hsa-miR-19b<br>hsa-miR-495<br>hsa-miR-92a | IGSF9B<br>SCN1A<br>LIFR<br>RGS4<br>EBF1<br>CSRNP3<br>COL19A1<br>OSBPL10<br>CFTR<br>CELSR2<br>CELF3<br>SYVN1<br>CTNND2<br>SERTAD2<br>GPR37L1<br>ADARB2<br>PDPK1 | hsa-miR-20a, hsa-let-7g<br>hsa-miR-98, hsa-let-7f, hsa-miR-93, hsa-miR-94, hsa-miR-95, hsa-miR-15b, hsa-miR-532-5p, hsa-miR-1, hsa-miR-146b-5p, hsa-miR-30e*, hsa-miR-142-5p<br>hsa-miR-153<br>hsa-miR-27a<br>hsa-miR-141<br>hsa-miR-374b* | DLST, CD969, ZNF652, PPP3R1, LZIC, DUSP8<br>RPS6KA3, PPP6R2, RAB12, DPH5, GBF1, ZF652, RPS6KA3, CELF1, L2HGDH, SUGT1, GIMAP, ZBTB16, GBP1<br>GBP3<br>TMSB4X<br>ABCA1<br>CRISPLD2<br>ACYP2<br>ACVR1<br>GBP2<br>SEC24A |
|  |  | NCAM2<br>WIF1<br>ANKRD7<br>SOX4<br>GDF11<br>TFDP2<br>EOMES<br>PLEKHA1<br>SLC4A10<br>SLC12A2<br>DLG5<br>CRTAM<br>PHLDA1 |  | EGR3<br>EGR2<br>CACNA1C<br>KIAA0319L<br>SLC7A14 |
| **SPMS** | none | none | hsa-miR-93, hsa-miR-15b, hsa-miR-20a,20b, hsa-let-7f,i,d,g, hsa-miR-97, hsa-miR-148b, hsa-miR-27b, hsa-miR-26b | PPP6R2, ZBTB16, DLST, NAA20, GIMAP7 DPH5 SNX8,LPAR6,GCNT1, MFHAS1,HINT1METAP2 |

### 2. Bioinformatic Analysis

#### 2.1. Gene-expression profile

The gene expression omnibus database (GEO, http://www.ncbi.nlm.nih.gov/geo) (Barrett *et al*., 2013), was used for mRNA and miRNA differential gene expression studies in multiple sclerosis samples from patients at different stages. GEO serves as a public repository for accessing high throughput data and microarray data (Edgar *et al*., 2002).

The GEO dataset accession numbers used in this study are provided on Github, https://github.com/omicscodeathon/ms_degs/tree/main/accessions

#### 2.2. Data analysis of differential expression of miRNAs and mRNAs

##### 2.2.1. GEO2R analysis

GEO2R is an online web-based tool for identifying differentially expressed genes. GEO2R analysis was performed on the microarray datasets retrieved from the GEO database (GSE21079, GSE17048 and GSE41890). This is done by assigning samples to a defined group based on their condition (RRMS, SPMS, PPMS and healthy control: HC) and also based on their sex. The differentially expressed mRNAs (DEG) and miRNAs were selected with a criteria of p<0.01 and |logFC|>0, while differentially expressed mRNA and miRNA genes with a criterion of p>0.01 were discarded using Benjamini and Hochberg’s False discovery rate (Benjamini and Yekutieli, 2001). Log transformation, limma precision weights (vooma) and force normalization were applied to the data.

##### 2.2.2. R analysis

GSE124900 study was analyzed using DESeq2, a widely used Bioconductor package in R for differential gene expression analysis of RNAseq datasets. This package employs the negative binomial distribution model to fit all genes across all samples and uses the Wald test to test for significance. DESeq2 internally normalizes the expression values by calculating the geometric mean. The gene count in each sample is then divided by the geometric means. The size factor for each sample is the median of the gene count-geometric mean ratio. This process corrects for bias in library size and RNA composition, which might occur when only a small number of genes are strongly expressed under one experimental condition but not under the other (Love *et al*., 2014). The differentially expressed miRNAs (DEMs) for this were selected with criteria of *padj <0.05* and |logFC|>0.

#### 2.3. miRNA-mRNA regulatory interaction

Mienturnet (Licursi *et al*., 2019), was used for predicting the target genes of the DEMs and for constructing the miRNA-mRNA regulatory relation. We determined the intersection between DEMs of miRNA in the datasets (GSE21079 and GSE125900) and the target DEGs of the mRNA datasets (GSE17048 and GSE41890). To say that the miRNA is a regulator of a gene, it must be antisense (miRNA downregulated and gene upregulated and vice versa).

#### 2.4. Gene Ontology and functional annotation of mRNA

The differentially expressed genes from microarray studies were first annotated by using the clariom package in R and therefore the genes were annotated with the Clariom D database package in R (MacDonald, 2021). After obtaining all the symbols for genes that were differentially expressed, Gene Ontology in R was performed using the ClusterProfiler package (Wu *et al*., 2021). Specifically, we used the enrichGO function in R, which also utilized the AnnotationDbi package (Pagès *et al*., 2022) and Org.Hs.eg.db package (Carlson, 2019). The output from Gene Ontology (GO) analysis was then converted to a data frame and plotted using the barplot function. Significant genes were considered based on *padj <0.05*.

## Results

### 1. Genes and miRNAs implicated in disease occurrence (RRMS VS HC)

#### 1.1. GSE124900 VS GSE41890

To determine the genes and miRNAs implicated in the disease onset, the first step aims to determine the differentially expressed miRNAs (DEM) and differentially expressed genes (DEGs) from the GSE124900 and GSE41890 datasets respectively.

The first study GSE124900 (N Baulina *et al*., 2019; Natalia Baulina *et al*., 2019) included 15 unrelated patients with RRMS (8 women and 7 men) and 8 healthy controls (4 women and 4 men) of comparable age. Among the 15 RRMS patients, 7 were in relapse and 8 were in remission. In this study, a total of 63 differentially expressed miRNAs (DEMs) (38 upregulated and 25 downregulated) were identified and distributed between women and men as follows: 8 RRMS women patients (4 in relapse and 4 in remission) and 4 HC. We observed 16 DEMs only between RRMS-rem and HC (8 up-regulated and 8 down-regulated) (Fig. 4A). While the comparison between 7 RRMS men patients (3 in relapse and 4 in remission) and 4 HC shows 5 DEMs between RRMS-rem and HC (2 up-regulated and 3 down-regulated), there were 42 DEMs between RRMS-rel and HC (28 up-regulated and 14 down-regulated) (Fig. 4B).

**Figure 4.**
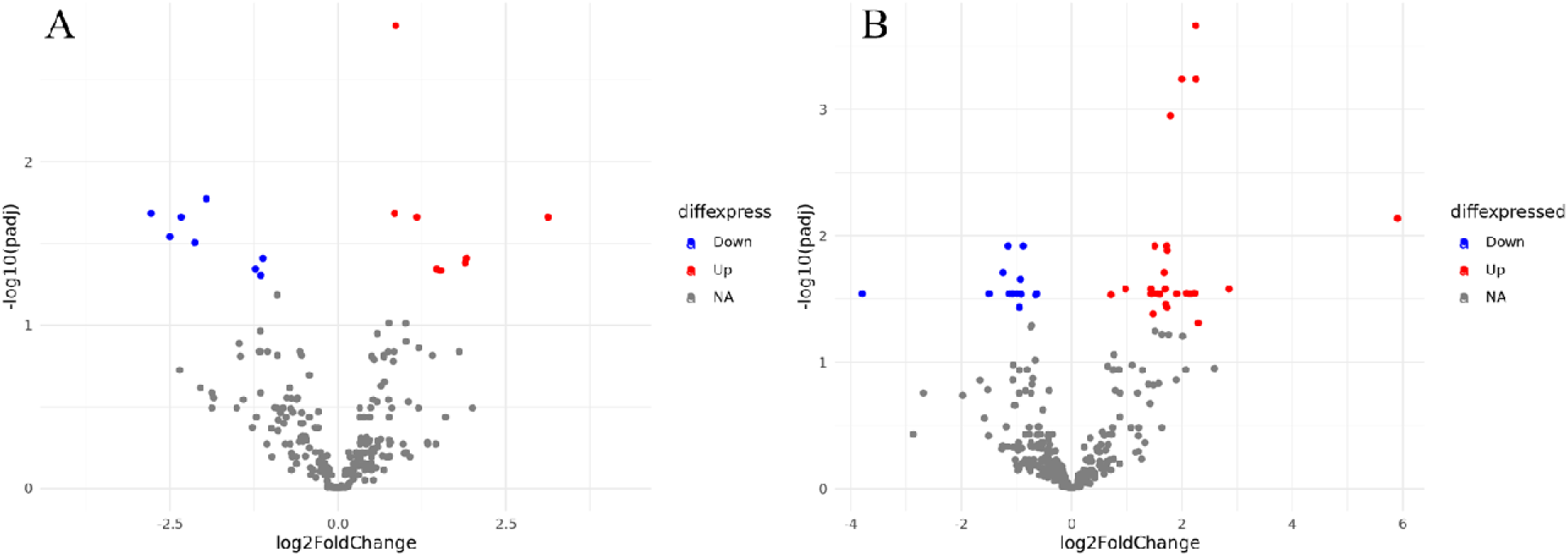
Expression profiles of DEMs. (A) Volcano plot of differentially expressed miRNA within women. The red dot represents upregulated miRNAs and the blue dot represents down-regulated miRNAs. (B) Volcano plot of differentially expressed miRNAs within men. The red dot represents upregulated miRNAs and the blue dot represents down-regulated miRNAs.

The GSE41890 dataset (H Irizar and Otaegui, 2013; Haritz Irizar *et al*., 2014) included 22 RRMS patients (12 women and 10 men) with samples both in remission and relapse cases (2 samples per patient) and 24 healthy controls (12 women and 12 men). After doing differential gene expression analysis using the GSE41890 study, we found 1,514 DEGs (733 upregulated and 781 downregulated) distributed between women and men. We analysed 12 RRMS women patient data (in relapse and remission) and 12 HC and noted 668 DEGs between RRMS-rel and HC (304 up-regulated and 364 down-regulated) (Fig. 5A). 223 DEGs were noted between RRMS-rem and HC (115 up-regulated and 108 down-regulated) (Fig. 5B). We noted 103 DEGs between RRMS-rel and RRMS-rem (40 up-regulated and 63 down-regulated) (Fig 5C). The comparison between 10 RRMS men patients (in relapse and remission) and 12 HC showed 420 DEGs between RRMS-rel and HC (211 up-regulated and 209 down-regulated), 72 DEGs between RRMS-rem and HC (57 up-regulated and 15 down-regulated) (Fig. 5D), and 28 DEGs between RRMS-rel and RRMS-rem (6 up-regulated and 22 down-regulated) (Fig. 5E).

**Figure 5.**
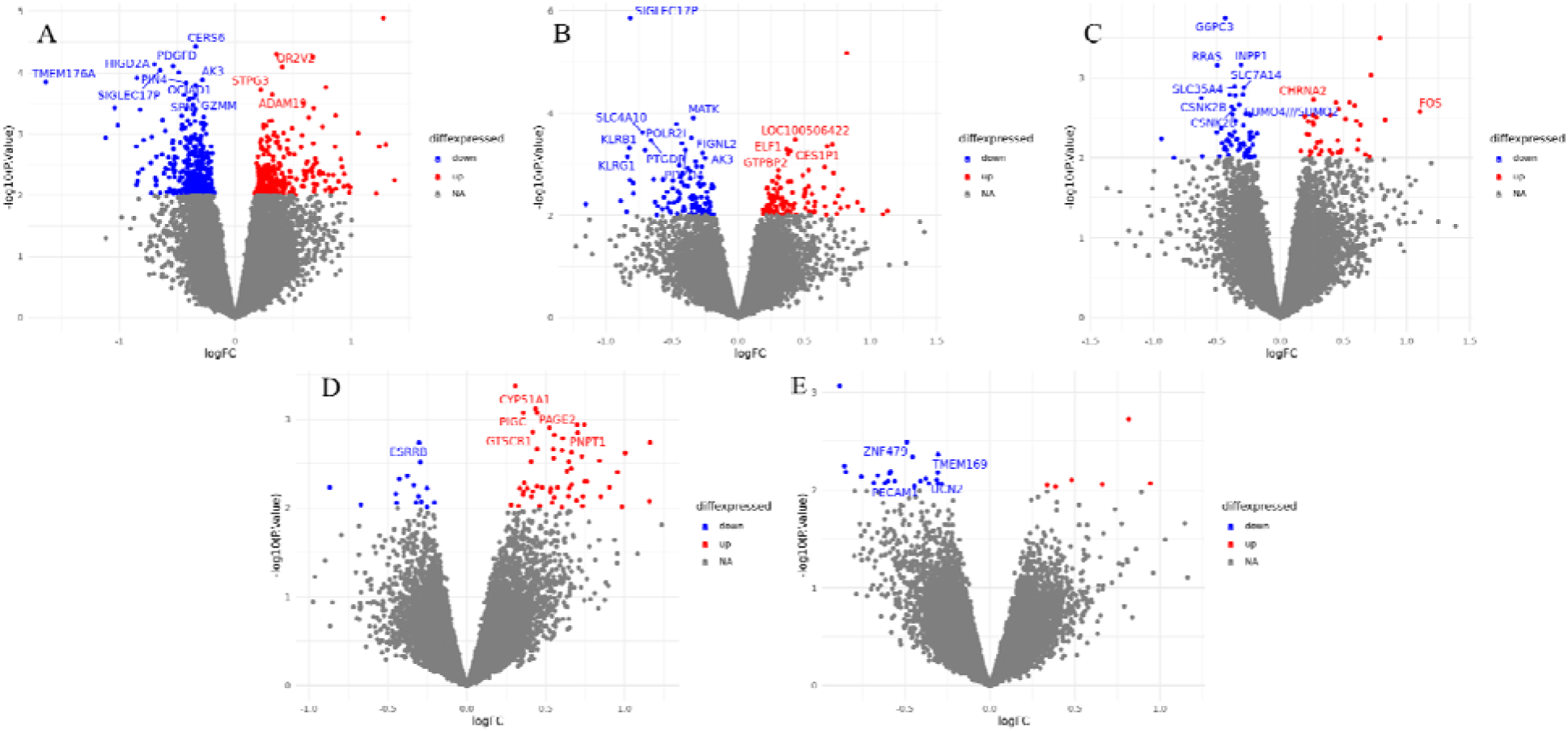
Expression profiles of DEG within women. The red dot represents upregulated mRNAs and the blue dot represents down-regulated mRNAs. (A) Volcano plot of differentially expressed mRNAs between RRMS-rel and HC. (B) Volcano plot of differentially expressed mRNAs between RRMS-rem and HC. (C) Volcano plot of differentially expressed mRNAs between RRMS-rel and RRMS-rem. (D) Volcano plot of differentially expressed mRNA between RRMS-rem and HC. (E) Volcano plot of differentially expressed mRNAs between RRMS-rel and RRMS-rem.

The second part of the study was the construction of the miRNA-mRNA regulatory relationship between the genes involved in GSE124900 and GSE41890 using Mienturnet (Licursi *et al*., 2019) based on the DEMs and DEGs found in the first part. To do this, the common genes between the two datasets should first be identified to compare the common groups. From the GSE124900 dataset, we noted some DEMs between women in remission and healthy control (RRMS-rem VS HC), men in remission and healthy controls (RRMS-rem vs HC) and men in relapse and healthy controls (RRMS-rel vs HC). We compared only these groups with those of GSE41890. To conclude that a miRNA is a regulator of a gene, it must be antisense (miRNA downregulated and gene upregulated and vice versa).

From the results (Supplementary Table 1), we noted that within women (RRMS-rem vs HC), 6 miRNAs identified from the GSE124900 dataset (1 upregulated and 5 downregulated) were regulators of 18 genes identified from the GSE41890 dataset (8 upregulated and 10 downregulated). Within men (RRMS-rel vs HC), 16 miRNAs identified from the GSE124900 dataset (12 upregulated and 4 downregulated) were regulators of 48 genes identified from the GSE41890 dataset (9 upregulated and 39 downregulated). In contrast, no intersection was found between miRNAs identified from the GSE124900 dataset and mRNAs identified from th GSE41890 dataset within men (RRMS-rem vs HC).

The third part of our study was to determine the DEG found after constructing the miRNA-mRNA regulatory relationship between the genes identified from the GSE124900 dataset and the GSE41890 dataset to find out the functions and processes of these genes associated with MS using the GO database. The results obtained showed that, within women (RRMS-rem vs HC), several significant DEGs are associated with multiple biological processes and molecular functions. However, no significant genes were found for cellular components neither for upregulated genes nor for downregulated genes (Fig. 6).

**Figure 6.**
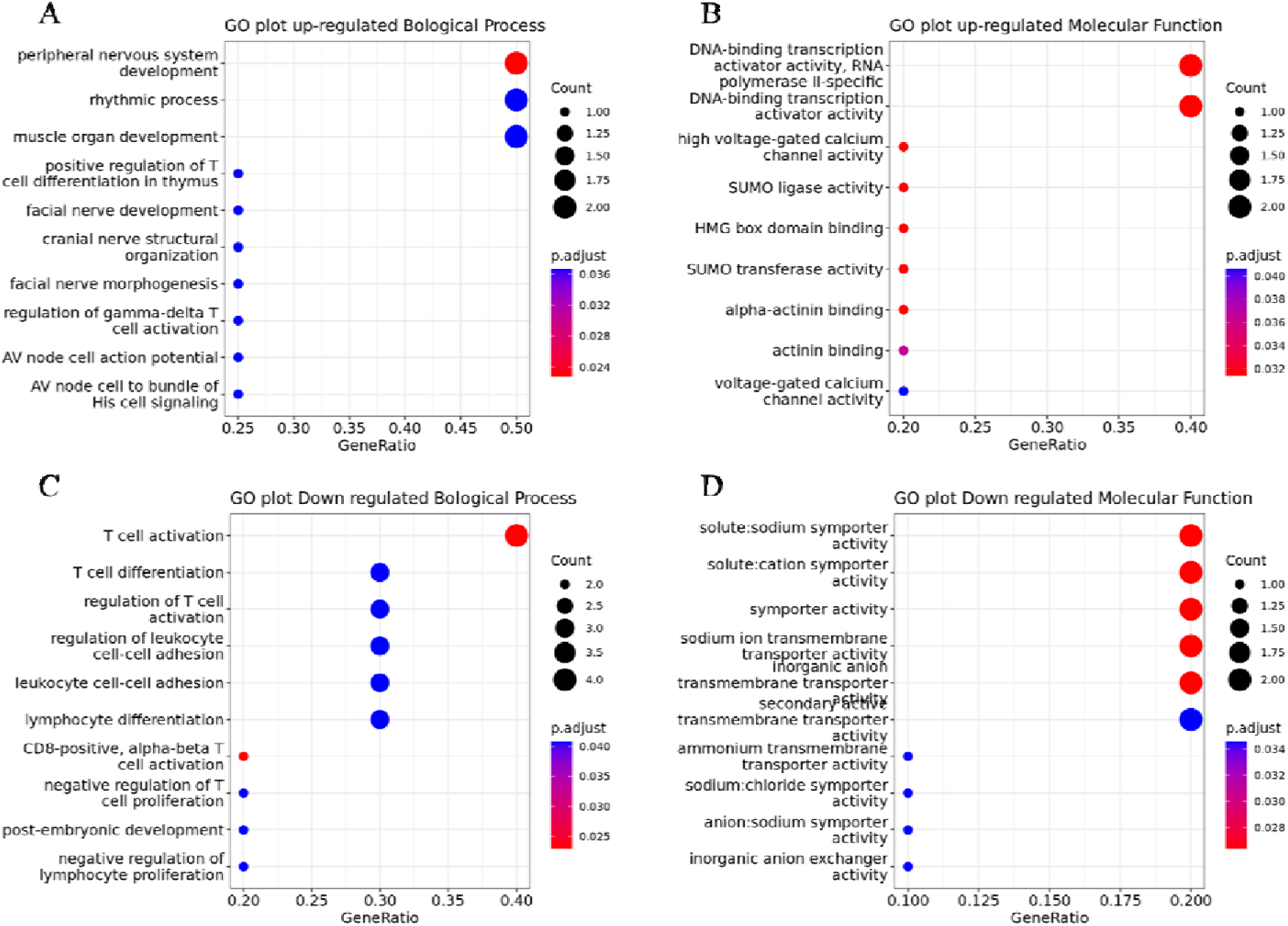
GO for DEGs within women RRMS-rem VS HC.

Within men (RRMS-rel vs HC), cellular response to interleukin-1 and cellular response to interferon-gamma are 2 of the biological processes where the upregulated genes are enriched (Fig. 7A). GTP binding and cytokine binding are molecular functions associated with the upregulated genes (Fig 7B). No significant genes were found for cellular components of upregulated genes. Downregulated genes do not present any significant gene for all the domains (biological process, molecular function and cellular component).

**Figure 7.**
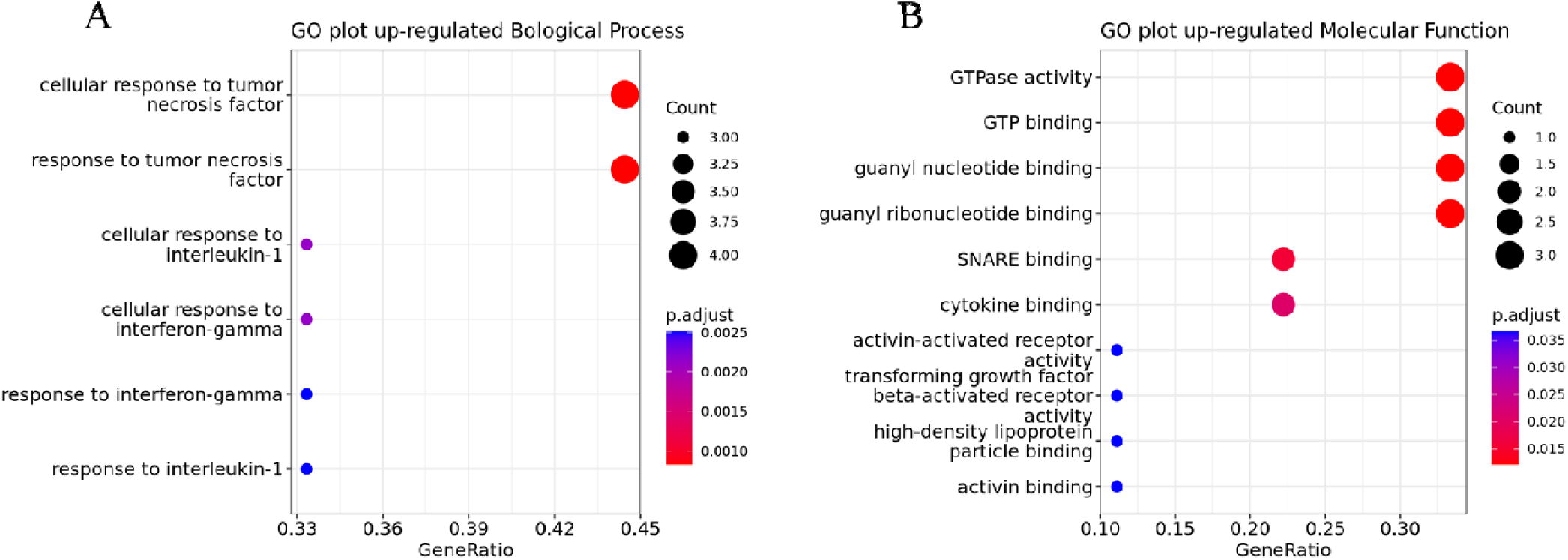
GO for DEG genes within men RRMS-rem vs HC

### 2. Genes and miRNAs implicated in disease progression (RRMS VS SPMS VS PPMS VS HC)

#### 2.1. GSE21079 VS GSE17048

To detect the mRNA genes and miRNAs implicated in the disease progression, the same steps that were followed in the determination of disease occurrence genes were followed. As a first step, we identify the differentially expressed miRNAs (DEMs) and differentially expressed genes (DEGs) from the studies GSE21079 and GSE17048 respectively. These two studies were carried out on an Australian population.

The first dataset GSE21079 (M B Cox *et al*., 2010) contains 59 MS patients and 37 HC. MS patients are divided into 24 RRMS patients (21 women and 3 men), 17 SPMS patients (13 women and 4 men) and 17 PPMS patients (6 women and 11 men). A total of 140 DEMs (47 upregulated and 93 downregulated) were identified from the dataset GSE21079. The volcano plots of the DEMs within women and men are depicted in Fig. 8 and Fig. 9 respectively. We combined the DEMs in both male and female groups and compared them with the DEGs identified from the GSE17048 dataset. From the GSE21079 dataset, we found 21 DEMs (17 downregulated and 4 upregulated) for RRMS vs HC, 45 DEMs (31 downregulated and 14 upregulated) for SPMS vs HC, 45 DEMs (28 downregulated and 17 upregulated) for PPMS vs HC, 10 DEMs (9 downregulated and 1 upregulated) for SPMS vs RRMS, 14 DEMs (5 downregulated and 9 upregulated) for PPMS vs RRMS and 5 DEMs (3 downregulated and 2 upregulated) for PPMS vs SPMS.

**Figure 8.**
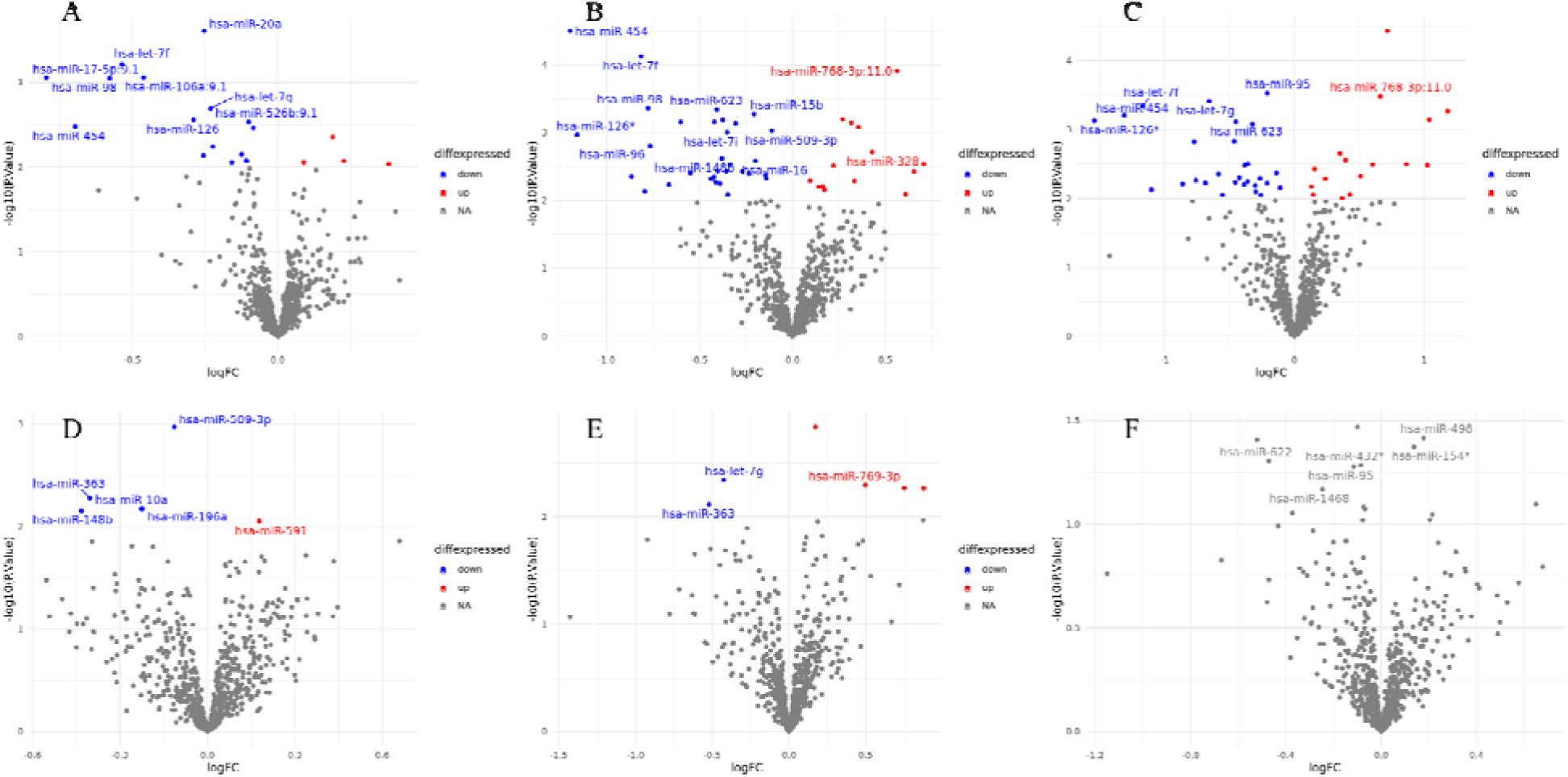
Expression profiles of DEM within women. The red dot represents upregulated miRNAs and the blue dot represents down-regulated miRNAs. (A) Volcano plot of differentially expressed miRNAs between RRMS and HC. (B) Volcano plot of differentially expressed miRNAs between SPMS and HC. (C) Volcano plot of differentially expressed miRNAs between PPMS and HC. (D) Volcano plot of differentially expressed miRNAs between SPMS and RRMS. (E) Volcano plot of differentially expressed miRNAs between PPMS and RRMS. (F) No differentially expressed miRNAs were noted between PPMS and SPMS.

**Figure 9.**
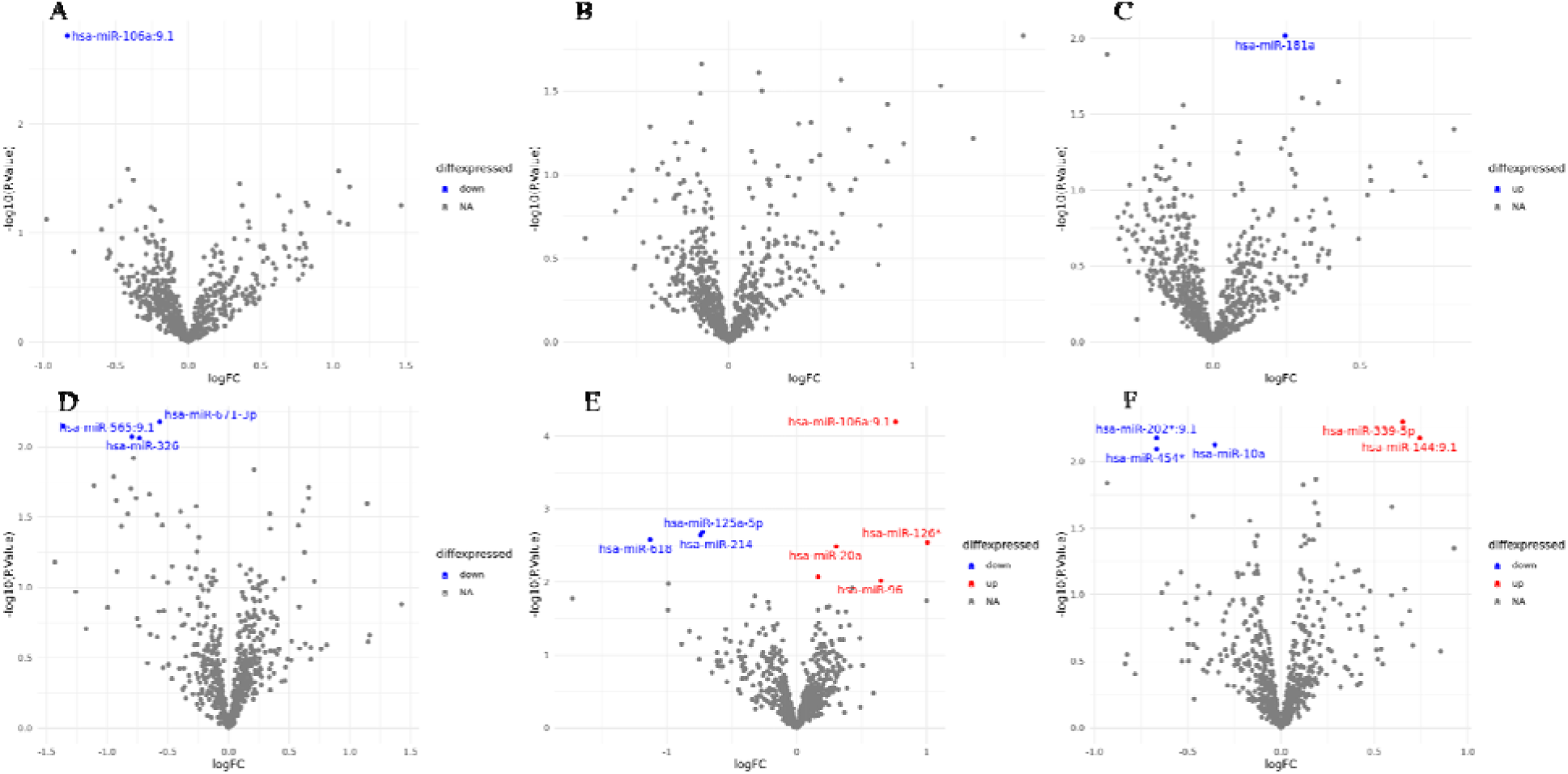
Expression profiles of DEMs within men. The red dot represents upregulated miRNAs and the blue dot represents down-regulated miRNAs. (A) Volcano plot of differentially expressed miRNAs between RRMS and HC. (B) No differentially expressed miRNAs between SPMS and HC. (C) Volcano plot of differentially expressed miRNAs between PPMS and HC. (D) Volcano plot of differentially expressed miRNAs between SPMS and RRMS. (E) Volcano plot of differentially expressed miRNAs between PPMS and RRMS. (F) No differentially expressed miRNAs were noted between PPMS and SPMS.

The GSE17048 dataset contains 99 MS patients (36 RRMS, 20 SPMS and 43 PPMS) and 45 healthy controls. A total of 1,404 DEGs were identified and distributed as follows: 422 DEGs (188 downregulated and 234 upregulated) for RRMS vs HC (Fig. 10A), 274 DEGs (146 downregulated and 128 upregulated) for SPMS vs HC (Fig. 10B), 581 DEG (296 downregulated and 285 upregulated) for PPMS vs HC (Fig. 10C), 29 DEGs (15 downregulated and 14 upregulated) for SPMS vs RRMS (Fig 10D), 68 DEGs (27 downregulated and 41 upregulated) for PPMS vs RRMS (Fig 10E) and 30 DEGs (10 downregulated and 20 upregulated) for PPMS vs SPMS (Fig 10F).

**Figure 10.**
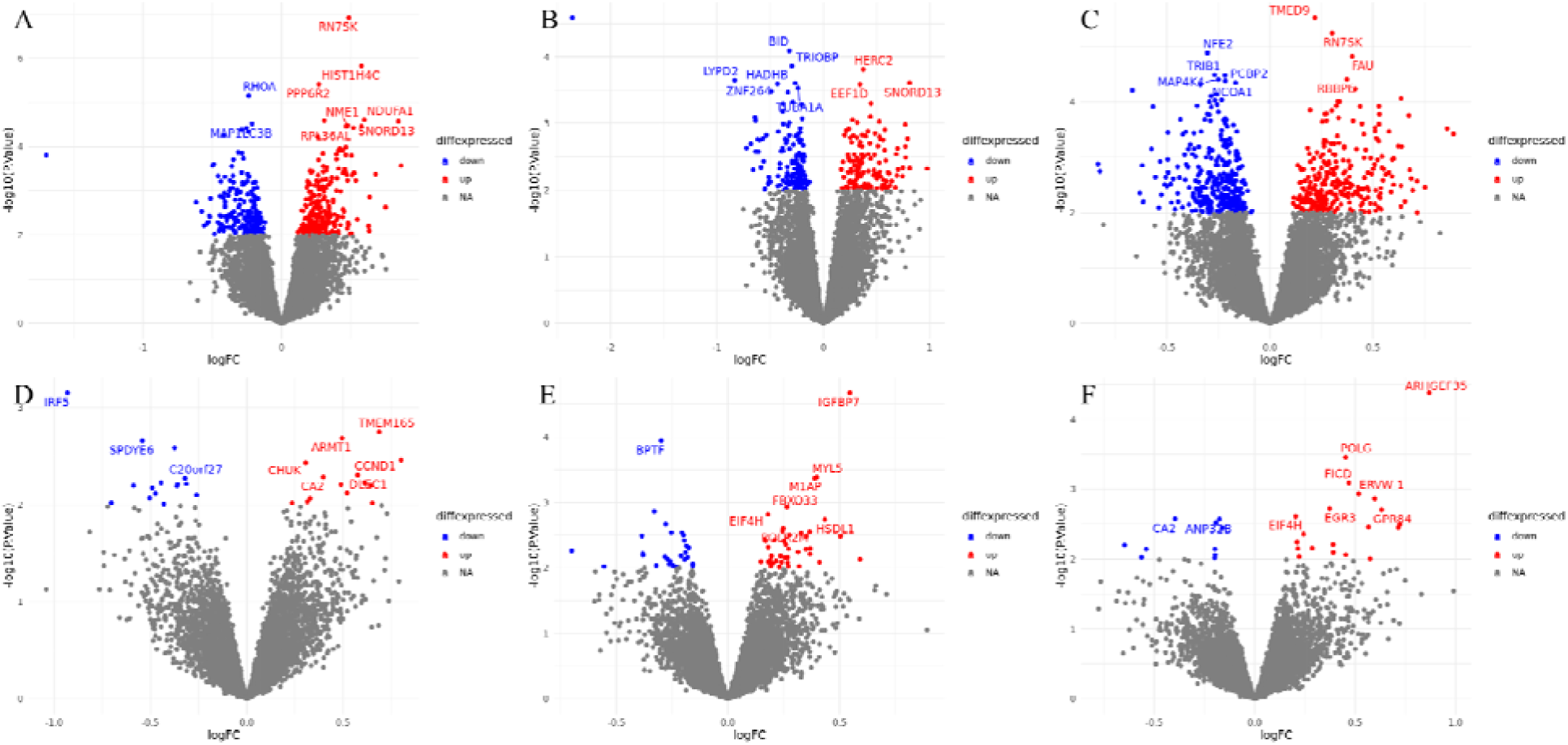
Expression profiles of DEG. The red dot represents upregulated mRNAs and the blue dot represents down-regulated mRNAs. (A) Volcano plot of differentially expressed mRNA between RRMS and HC. (B) Volcano plot of differentially expressed mRNAs between SPMS and HC. (C) Volcano plot of differentially expressed mRNAs between PPMS and HC. (D) Volcano plot of differentially expressed mRNAs between SPMS and RRMS. (E) Volcano plot of differentially expressed mRNAs between PPMS and RRMS. (F) No differentially expressed mRNAs were noted between PPMS and SPMS.

All the differentially expressed miRNAs and mRNAs are provided in our GitHub repository, (https://github.com/omicscodeathon/ms_degs/tree/main/output/TablesDEGs).

As a second step of the study, the miRNA target enrichment analysis was done using mienturnet to construct the miRNA-mRNA regulatory relationship between genes in the GSE21079 dataset and the GSE17048 dataset based on DEMs and DEGs found in the first step. The results show that between RRMS vs HC and SPMS vs HC groups, some of the upregulated and downregulated DEMs were found to target the downregulated and upregulated DEGs respectively as depicted in the supplementary Table 3, while the other groups show that there is no miRNA target activity between the DEMs and DEGs. In the final step, gene ontology was also performed for DEGs found after the construction of the miRNA-mRNA regulatory relationship between genes in the GSE21079 dataset and the GSE17048 dataset to determine the functions and processes of these genes associated with MS. Within the first group, RRMS vs HC, a single function was assigned to upregulated genes which is 2-oxoglutarate metabolic process as a biological process (Fig 11A). No significant function was assigned to the upregulated gene neither for molecular function nor for cellular function. The same result was observed for the downregulated genes for all the domains (biological process, molecular function and cellular component). Within the second group, SPMS vs HC, two functions were assigned to the upregulated genes which are peptidyl-methionine modification and N-terminal protein amino acid modification as biological processes (Fig. 11B). No significant function was assigned to the upregulated genes neither for molecular function nor for cellular function. The same result was observed for the downregulated genes for all the domains (biological process, molecular function and cellular component).

**Figure 11.**
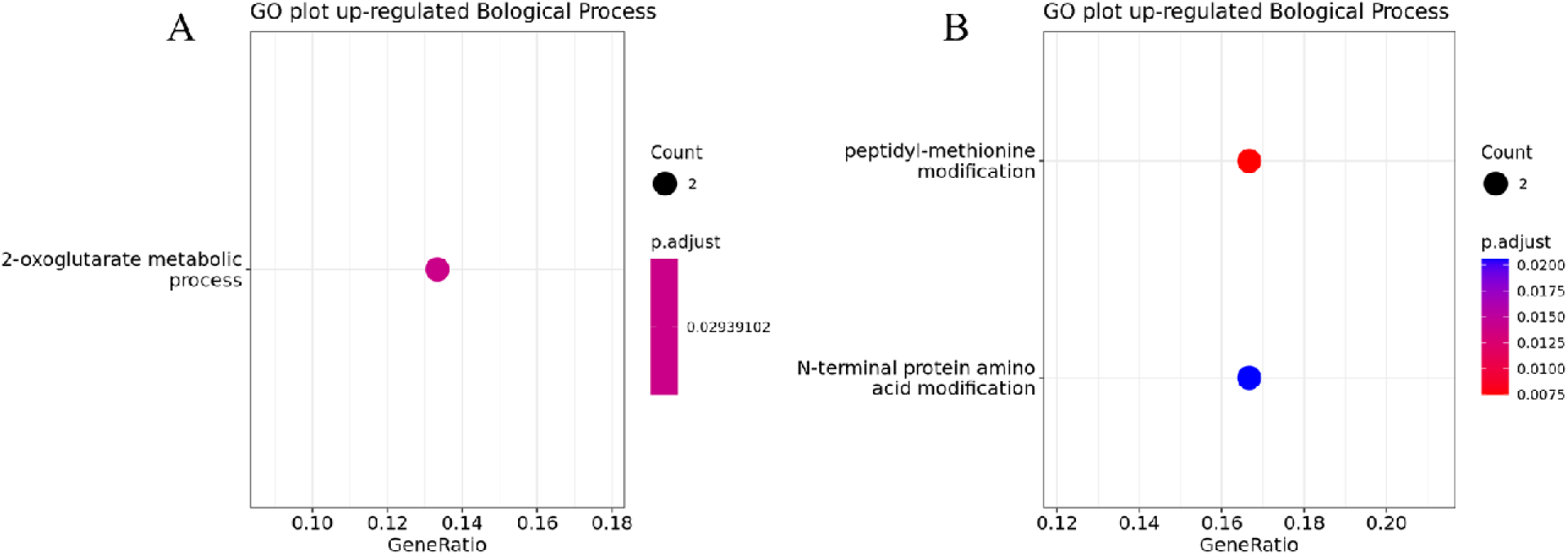
(A) GO for DEGs within RRMS vs HC group. (B) GO for DEGs within the SPMS vs HC group.

### 3. Common genes and miRNAs implicated in disease occurrence in different populations (RRMS VS HC)

#### 3.1. GSE21079 VS GSE124900

There are no common miRNAs between genes identified from the GSE21079 and GSE124900 datasets.

#### 3.2. GSE17048 VS GSE41890

There are 98 common mRNAs between genes identified from the GSE17048 and GSE41890 datasets, 57 of which are down-regulated in both datasets, while 41 are up-regulated in both. A full list of mRNAs is in supplementary table 4. Gene ontology was performed for all these common mRNAs. The results indicate the assigned activities for both upregulated and downregulated genes. For upregulated genes, RNA splicing is one of the biological processes that we noted (Fig. 12A). Furthermore, only one molecular function was assigned to these genes which is: the structural constituent of ribosome (Fig. 12B). In addition, several cellular component functions were related to the upregulated genes (Fig. 12C).

**Figure 12.**
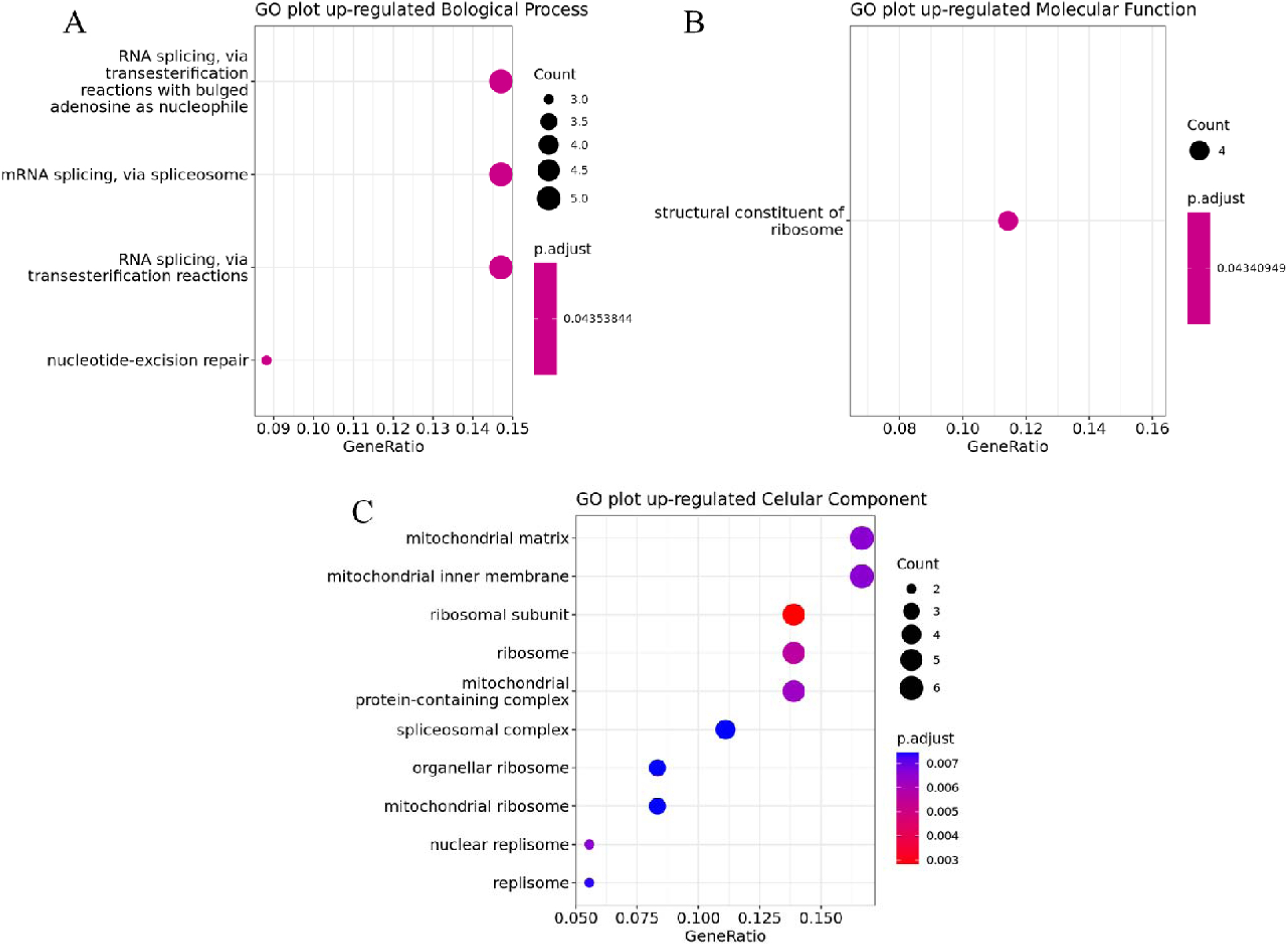
GO for common upregulated genes identified from the GSE17048 and GSE41890 datasets.

From the downregulated genes, two biological processes were noted: intracellular receptor signalling pathway and negative regulation of defence response to the virus (Fig. 13A). Moreover myosin binding is one of the molecular functions assigned to the latter (Fig. 13B). We noted, several cellular component functions that were related to these genes (Fig. 13C).

**Figure 13.**
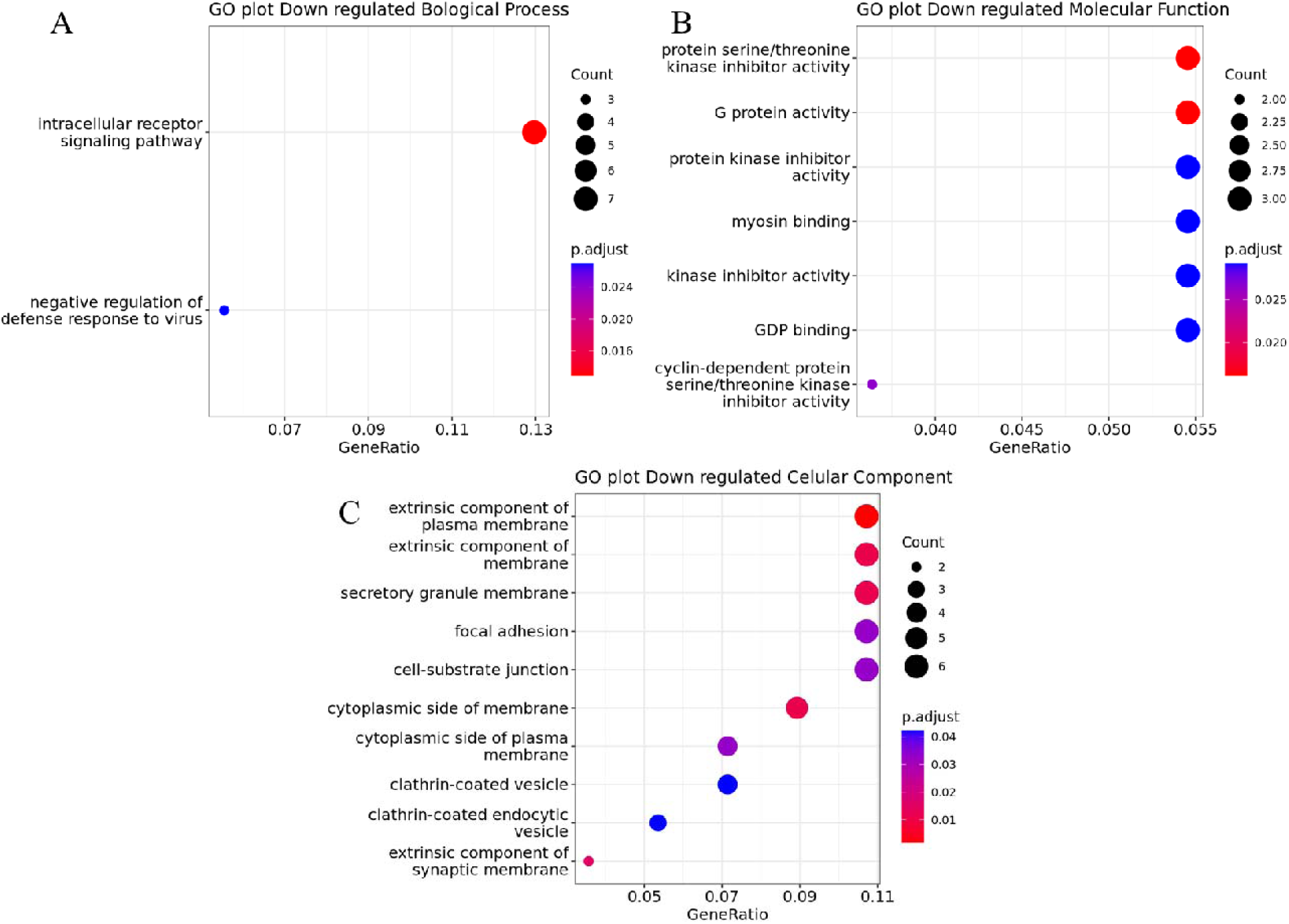
GO for common downregulated genes identified from the GSE17048 and GSE41890 datasets.

## Discussion

Multiple sclerosis is an autoimmune disease which occurs as a result of the demyelination of nerve cells in the central nervous system and it exists in 3 stages which are RRMS, SPMS and PPMS. This has made the diagnosis of a particular stage of MS difficult. Many methods that have been employed to diagnose the stages of MS depend on medical history and neurological examination with Magnetic Resonance Imaging (MRI) and cerebrospinal fluid analysis but they are not that accurate (Ömerhoca, Akkaş, and İçen, 2018). Therefore our study aims at identifying biomarkers specific to each stage of MS using a computational approach. These biomarkers could also be therapeutic target points. Here, we look at different GEO datasets to enhance our confidence across different populations, and our reports for each are therefore discussed.

We investigated the genes and miRNAs that are implicated in disease occurrence (RRMS vs HC). After we analysed the datasets GSE124900 and GSE41890, we noted that there were 1,514 DEGs of which 733 were found to be upregulated and 781 were found to be downregulated. There is a marked difference in the genes controlling the biological processes that are affected in women and men with RRMS (Fig. 6 and Fig. 7). As seen in Figure 6, many of the genes which are upregulated in women with RRMS are associated with the peripheral nervous system, rhythmic process and muscle organ development. However, in men, the upregulated genes affect the cellular response to tumour necrosis factor. In a previous study, it was found that the potential gene targets of specific upregulated miRNAs affect chemokines, transcription factors and cytokines like the tumour necrosis factor as well as growth factors and cell apoptosis regulators (Ma *et al*., 2014). The miRNA-146b was one of the specific miRNAs in Ma *et al*., (2014) affecting these pathways. However, it was found in our study to be downregulated but still with an effect of upregulation on other target genes such as ACYP2.

There were several biological processes affected by the upregulation of many target genes as a result of the downregulation of specific miRNAs in women with RRMS. These target genes were specifically associated with T-cell activation, differentiation and regulation as well as leukocyte cell-cell activation, adhesion and differentiation. In this study, results showed that miR-142-5p was under-expressed in the serum of RRMS-remission women patients compared to healthy controls. Talebi *et al*., (2017) evaluated the expression levels of miR142 in the brain of mice with experimental autoimmune encephalomyelitis (EAE: animal model of MS) and showed that the expression was increased in mouse models compared to controls. They also proved that both miR-142 isoforms influence the T differentiation leading to the autoimmune neuroinflammation. Also, miR-142-5p was reported to be highly expressed in the serum of patients (Junker *et al*., 2009).

We investigated the genes and miRNAs that are implicated in disease progression (SPMS vs PPMS). After we analysed the datasets GSE21079 and GSE17048, we noted that the proposed method in comparing samples (both the miRNA and mRNA) at different stages of multiple sclerosis (PPMS, SPMS, RRMS) and HC with mienturnet, identified 12 miRNAs implicated in SPMS and 6 implicated miRNAs have been linked to PPMS (Otaegui *et al*., 2009).

After analyzing the GSE21079 dataset of SPMS patients, we found that miRNA hsa-let-7f is downregulated in the patients as reported by (Li *et al*., 2019) with a lack of functionality to suppress Th17 cell differentiation. This is a key factor in the pathogenesis of MS and has also been shown to lead to overexpression of the dihydrolipoamide S-succinyltransferase gene (DSLT), which has been implicated in relapsing multiple sclerosis (La Rocca *et al*., 2017). Another study has shown that the downregulation of hsa-miR-15 and hsa-miR-20 has led to the upregulation of CD69, a leukocyte activation marker that has been involved in the pathogenesis of chronic inflammation (Sancho Gómez, and Sánchez-Madrid, 2005). However recent studies have reported it to play a controversial role in suppressing the differentiation of Th17 (González *et al*., 2013). The distinctive deregulation of the expression of hsa-miR-96 leads to the progression of RRMS (Sedeeq *et al*., 2019) and the distinctive deregulation of the expression of hsa-miR-27b leads to the progression of RRMS (Shafiei *et al*., 2020). From our study, we also found that hsa-miR-96 and hsa-miR-27b also play significant roles in the progression of SPMS. Furthermore, after analysing the GSE21079 dataset of RRMS patients, we noted a distinctive miRNA (hsa-miR-93) from the SPMS patients that were deregulated and over-expressed in a study by Shafiei *et al*., (2020). In this study, it was proposed to be a prospective biomarker. However, from our study, its deregulation has been shown to increase the expression of protein phosphatase 6 regulatory subunit 2 (PPP6R2), a scaffolding PP6 subunit which is involved in PP6-mediated dephosphorylation of NFKBIE, opposing its degradation in response to TNF-∝. This further validates our result with the study that shows that TNF-∝ is a major cytokine that contributes to the pathogenesis of MS (Zahid *et al*., 2021).

The gene ontology analysis of the upregulated target genes and the downregulated miRNAs of SPMS and RRMS from this study shows that all the genes contribute towards the pathogenesis of these stages of MS. This includes promoting the ribonucleoprotein biogenesis in terms of their biological process, acting as structural constituent of ribosomes in term of their molecular functions and also forming part of a membrane protein in term of their cellular function, which is explained in part by International Multiple Sclerosis Genetics Consortium, 2019. This helps to further validate our results that these downregulated miRNAs and upregulated mRNA are drivers that trigger multiple sclerosis to the various stages and can therefore be used as biomarkers for its representation. However, further analyses via the *in-vitro* techniques and pan-genomic-wide association studies need to be conducted on MS patients to further validate these biomarkers as representatives of each stage.

We investigated the common genes and miRNAs implicated in disease occurrence in different populations (RRMS vs HC). After we analysed the datasets GSE17048 and GSE41890, we noted that the GSE17048 study consists of 36 RRMS patients and the GSE41890 study consists of 22 RRMS patients. When comparing the GSE17048 and GSE41890 datasets, there were 98 common mRNAs in RRMS, 41 of which were up-regulated in both studies and 57 which were down-regulated in both studies. GO analysis of the up-regulated mRNAs shows that several genes are involved in RNA splicing and are structural constituents of the ribosome. In addition, many of the genes are found at various ribosomal sites. Defective RNA processing has previously been implicated in RRMS, particularly regarding structural RNA (LaCava *et al*., 2005). The processing of RNA is a vital step in gene expression. Polyadenylation of mRNAs, for instance, is necessary in the synthesis and maturation of mRNA. Noncoding RNAs lack the 3’ poly(A)tails. However, the addition of these can be used to promote the degradation of misfolded or incorrectly processed noncoding RNAs (Spurlock *et al*., 2015). Therefore, it is possible that the dysregulation of these processes could be a molecular mechanism underlying MS progression. Indeed, Spurlock *et al*., (2015) found that mononuclear cells from patients with RRMS exhibited increased levels of polyadenylated non-coding RNAs (LaCava *et al*., 2005). Furthermore, the treatment of these cells with a common MS therapy can restore the levels of some of these RNAs, suggesting that the dysregulated polyadenylation may have pathogenic consequences (LaCava *et al*., 2005). A study by Hao *et al*., (2022) also showed that hub genes targeted by long non-coding RNAs (lncRNAs) and competing endogenous RNAs (ceRNAs) functioned in ribosomal processes. In addition, gene set enrichment analysis also showed that highly expressed hub genes were associated with base excision repair. Indeed, from our analysis, nucleotide excision repair was also shown to be upregulated in our analysis. Interestingly, there have been reports of MS relapse in response to mRNA CoVID-19 vaccines. Toljan *et al*., (2022) concluded that acute neurological deficits in response to recent mRNA vaccine administration may represent a new onset of MS. While this association cannot be definitively determined to be causal, it does show that RNA changes may underlie the onset and progression of MS. Therefore, the dysregulation of RNAs and excision repair may be important mechanisms in RRMS. Interestingly, several genes are also found in the mitochondrial matrix and inner mitochondrial membrane. Mitochondrial dysfunction is well known to play a role in MS as well as other neurodegenerative disorders (Barcelos *et al*., 2019). Therefore, it is also possible that defective mitochondrial RNA processing could be implicated in MS progression.

GO analysis of the down-regulated mRNAs showed that there is less serine/ threonine kinase inhibition. Serine/ threonine kinases are important for the regulation of apoptosis (Cross *et al*., 2000). Therefore, if inhibitors of serine/ threonine kinases are down-regulated, this may hint at a molecular mechanism for the neuronal cell death observed in RRMS. We also observed decreased intracellular signalling and G protein activity, which work together to transmit signals to the cell surface. In agreement with this, several proteins are found in membranes and junctions of the cells, where signalling generally occurs. These signalling pathways often target transcription factors that regulate gene expression, thereby relating to the dysregulated RNA processing previously discussed. Interestingly, several mRNAs are also found in vesicles and at the synaptic membrane. This could show that decreased intracellular signalling and vesicle transport at the synapses may play a role in RRMS. Indeed, there is growing evidence that synapse dysfunction plays an important role in MS and that synaptic structures play a key role in disease pathophysiology (Bellingacci *et al*., 2021; Centonze *et al*., 2010).

Overall, for RRMS we found 30 miRNAs (13 upregulated and 17 downregulated) that could be potential biomarkers for RRMS specifically. Similarly, for SPMS we found 9 downregulated miRNAs as potential biomarkers. No potential biomarkers for PPMS were found in this study. These miRNAs are involved in diverse roles in MS and could potentially be used in conjunction to improve their efficacy. Further work would need to be done on the specific miRNAs found, but, once validated, they could be used as biomarkers for disease occurrence and progression (Supplementary table 5).

## Conclusion

In the overall analysis of MS in this study, we were able to report some biomarkers that would likely be found in RRMS and SPMS patients, but we could not report any for PPMS during the screening process. However, by comparing multiple studies we have been able to determine which mRNA genes and miRNAs should be studied in more detail. Therefore further analysis via *in-vitro* techniques and pangenomic-wide association studies needs to be conducted on multiple sclerosis patients to further validate these biomarkers and those yet unidentified as representative of each stage of MS. Once validated, these miRNAs and mRNAs may assist clinical and therapeutic decisions and improve treatment outcomes.

## Availability and Requirements

Project name: Multiple Sclerosis Stages and their Differentially Expressed Genes: A Bioinformatics Analysis

Project home page: https://github.com/omicscodeathon/ms_degs

## Abbreviations

CNS: Central Nervous System
DEG: Differentially Expressed Genes
DEM: Differentially Expressed miRNAs
EAE: experimental autoimmune encephalomyelitis
GEO: Gene Expression Omnibus
GO: Gene ontology
HC: Healthy control
miRNA: micro ribonucleic acid
mRNA: Messenger ribonucleic acid
MS: Multiple sclerosis
NCBI: National Center for Biotechnology Information
PPMS: Primary Progressive Multiple Sclerosis
RRMS: Relapsing-Remitting Multiple sclerosis
SPMS: Secondary Progressive Multiple Sclerosis

## Acknowledgements

The authors thank the National Institutes of Health (NIH) Office of Data Science Strategy (ODSS) and the Genetics Society of America for their immense support before and during the October 2022 Omics codeathon organized in collaboration with the African Society for Bioinformatics and Computational Biology (ASBCB).

## Author Contributions

FA and GB conceived the original idea. FA, IA, MTK and DAA developed the bioinformatics analysis pipeline. IA, MTK and DA performed the bioinformatics analysis of the case study data and FA helped in some analysis. All authors validated the pipeline. GB and FA interpreted the results and provided feedback. FA, KC, DAA and NS drafted the manuscript. OIA reviewed the manuscript, provided critical feedback and helped shape the final version of the manuscript. OIA’s role was in the administration and supervision of the bioinformatics analysis in the project. OIA also provided the resources to facilitate and complete the analysis and provided guidance, editing and final review of the manuscript. All authors read and approved the final manuscript.

## Declarations

### Ethics approval and consent to participate

Not applicable.

### Consent for publication

Not applicable.

